# MED1 is a lipogenesis coactivator required for postnatal adipose expansion

**DOI:** 10.1101/2020.11.16.384867

**Authors:** Younghoon Jang, Young-Kwon Park, Ji-Eun Lee, Nhien Tran, Oksana Gavrilova, Kai Ge

## Abstract

MED1 often serves as a surrogate of the general transcription coactivator complex Mediator for identifying active enhancers. MED1 is required for phenotypic conversion of fibroblasts to adipocytes *in vitro* but its role in adipose development and expansion *in vivo* has not been reported. Here we report that MED1 is dispensable for adipose development in mice. Instead, MED1 is required for postnatal adipose expansion and the induction of *de novo* lipogenesis (DNL) genes after pups switch diet from high-fat maternal milk to carbohydrate-based chow. During adipogenesis, MED1 is dispensable for induction of lineage-determining transcription factors (TFs) PPARγ and C/EBPα but is required for lipid accumulation in the late phase of differentiation. Mechanistically, MED1 controls the induction of DNL genes by facilitating lipogenic TF ChREBP-dependent recruitment of Mediator to active enhancers. Together, our findings identify a cell- and gene-specific regulatory role of MED1 as a lipogenesis coactivator required for postnatal adipose expansion.

- MED1 is largely dispensable for adipogenesis and embryonic development of adipose tissue
- MED1 is required for postnatal adipose expansion
- MED1 is required for DNL gene expression in adipocytes
- MED1 controls DNL gene transcription by facilitating ChREBP-dependent Mediator binding to active enhancers

## INTRODUCTION

Adipose tissue development starts before birth. After birth, adipose tissues, especially white adipose tissue (WAT), undergo marked expansion with tissue mass and lipid contents increasing from newborn pups to adult mice^1,2^. Adipose tissue development requires the master adipogenic transcription factor (TF) PPARγ, which is induced in the early phase of adipogenesis, the differentiation of preadipocytes towards adipocytes^3^. PPARγ cooperates with C/EBPα on transcriptional enhancers to induce the expression of thousands of adipocyte genes that also include major lipogenic TFs ChREBP and SREBP1, which together promote *de novo* lipogenesis (DNL) in the late phase of adipogenesis^4,5^.

Liver and adipose tissues are the main sites for DNL in mammals. Fatty acid synthesis is low in liver and adipose tissue of mouse pups but increases dramatically after the switch of diet from high-fat maternal milk to carbohydrate-rich standard laboratory chow^6^. DNL is a tightly regulated metabolic process that converts excess carbohydrates to fatty acids before the synthesis of triglyceride for energy storage. Through glycolysis and the tricarboxylic acid cycle, carbohydrates are converted to citrate. Fatty acid synthesis enzymes, including ATP citrate lyase (ACLY), acetyl-CoA carboxylase α/β (ACACA/ACACB, also known as ACC1/ACC2), fatty acid synthase (FASN) and stearoyl-CoA desaturase-1/2/3/4 (SCD1/2/3/4), act sequentially to convert citrate to fatty acids^7,8^. Through the subsequent actions of triglyceride synthesis enzymes GPAM (also known as GPAT1), 1-acylglycerol-3-phosphate O-acyltransferase-1/2 (AGPAT1/2), LIPIN1 and DGAT1, fatty acids are esterified with glycerol-3-phosphate and incorporated into triglyceride^9^.

DNL enzyme expression is transcriptionally regulated by major lipogenic TFs carbohydrate-responsive element-binding protein (ChREBP, also known as MLXIPL) and sterol regulatory elementbinding protein 1 (SREBP1) in adipose tissues. ChREBP is a major driver for adipocyte DNL^8^. It is expressed as two isoforms ChREBPα and ChREBPβ, which are encoded by the same gene but transcribed from different promoters. ChREBPβ transcription activity is ~20-fold higher than that of ChREBPα^10^. ChREBPα is constitutively expressed but is sequestered in the cytosol and transcriptionally inactive under low glucose conditions. ChREBPα is activated by high glucose and, in turn, stimulates ChREBPβ expression^11^. Overexpression of the constitutively active form of ChREBP (CA-ChREBP) in mouse WAT induces expression of DNL enzymes such as ACLY, ACACA, FASN and SCD1^12^. Conversely, adipocyte-specific knockout (KO) of ChREBPα/β reduces the expression of ACLY, ACACA, FASN and SCD1 and abrogates glucose-induced DNL in WAT^13^. ChlP-Seq analysis in mouse WAT showed genomic binding of ChREBP on *Acaca, Acacb, Fasn* and *Chrebp* genes suggesting that ChREBP directly regulates transcription of DNL enzymes as well as itself^14^. SREBP1 seems to play a minor role in DNL in adipocytes. While Fabp4-promoter driven overexpression of the SREBP1a isoform increases ACACA and FASN expression in WAT and BAT^15^, whole body KO of SREBP1 does not affect adiposity or DNL enzyme expression in WAT^16^.

In eukaryotes, all transcription is mediated by RNA Polymerase II (Pol ll) for protein-coding genes. Transcription by Pol ll requires transcriptional co-regulators including chromatin remodelers, histone modifiers and the Mediator coactivator complex^17^. The Mediator complex comprises approximately 30 subunits and it generally serves as a functional bridge between TFs and basal transcriptional machinery including Pol II^18^. Mediator is enriched at transcriptional promoters and enhancers. One of the Mediator subunits, MED1 (also known as TRAP220, PBP), is often used as a surrogate for the Mediator to identify active enhancers and super-enhancers^19^. Whole-body *Med1* KO in mice leads to embryonic lethality around E1 1.5^20,21^.

We previously reported that *Med1* KO mouse embryonic fibroblasts show defects in PPARγ- and C/EBPβ-stimulated phenotypic conversion into adipocytes^22,23^. However, it has remained unclear whether MED1 is required for adipose tissue development and expansion *in vivo.* Using Adipoq-Cre- or Myf5-Cre-mediated deletion of *Med1* in adipocytes or precursor cells in mice, we show that MED1 is required for postnatal adipose expansion after switching diet from high-fat maternal milk to carbohydrate-based standard chow and that Med1 is largely dispensable for adipose tissue development. By RNA-Seq and ChIP-Seq analyses, we found that MED1 directly controls the transcription of DNL genes including *Acly, Acaca, Scd1* and *Fasn* in adipocytes, but not the expression of early adipogenesis marker *Pparg.* MED1 facilitates Mediator binding on active enhancers of ChREBP-regulated lipogenesis genes in adipocytes.

## RESULTS

### *Med1*^f/f^;*Adipoq-Cre* mice show lipodystrophy and are resistant to high fat diet-induced obesity

To investigate the functional role of MED1 in adipose tissues, we used conditional knockout (KO) mice targeting exons 8-10 of *Med1* gene (*Med1*^f/f^, hereafter referred to as f/f) (Figure S1A)^24^. Adipocyte-specific *Med1* KO *(Med1^f/f^;Adipoq-Cre,* A-KO) mice were generated by crossing f/f mice with *Adipoq-Cre* mice expressing *Adiponectin* promoter-driven Cre^25^. A-KO mice showed similar appearance and total body weight compared to f/f mice but had significantly reduced fat mass and increased lean mass at 8 weeks (8wk) (Figure 1A-B). Adipose tissues including BAT, inguinal white adipose tissue (iWAT) and epididymal WAT (eWAT) decreased in A-KO mice, while liver mass increased (Figure 1C-D). Similar results were obtained from female mice at 8wk (Figure S1B-E). Histological analysis revealed reduced lipid droplets in adipose tissues of A-KO mice (Figure 1E). Deletion of *Med1* was verified in adipose tissues (Figure 1F). These results indicate that deletion of *Med1* in adipocytes leads to lipodystrophy in adult mice.

**Figure 1.**
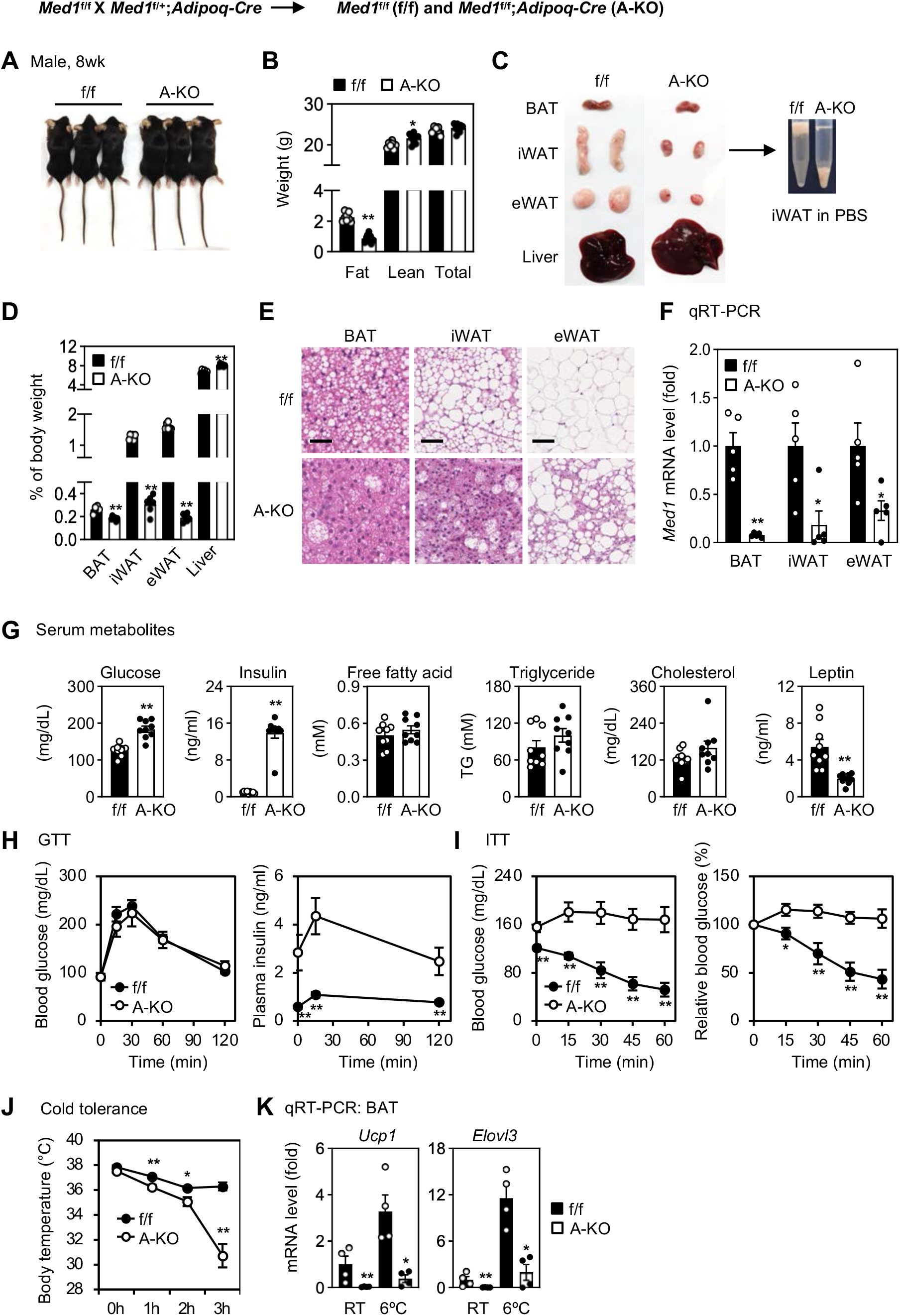
Deletion of *Med1* in adipocytes leads to lipodystrophy. (A-F) Deletion of *Med1* in adipocytes leads to lipodystrophy in adult mice. *Med1*^f/f^ (f/f) mice were crossed with *Adipoq-Cre* to generate adipocyte-specific *Med1* KO mice (A-KO). All data were from 8 weeks (8wk) male mice fed with a regular diet. Data from female mice and metabolic studies are shown in Figure S2-3. (A) Representative morphology of adult mice. (B) Body composition. Fat mass, lean mass and total body weight were measured by MRI (*n*=8 per group). (C) Representative pictures of brown adipose tissue (BAT), inguinal white adipose tissue (iWAT), epididymal WAT (eWAT) and liver. iWAT from A-KO mice failed to float in PBS. (D) Average tissue weights are presented as % of body weight (*n*=6 per group). (E) H&E staining shows reduced lipid droplet in A-KO adipose tissues. *Scale bar =* 100 μm. (F) qRT-PCR analysis of *Med1* mRNA levels in BAT, iWAT and eWAT (*n*=5 per group). (G) Levels of serum metabolites (*n*=8 per group). (H) Glucose tolerance test (GTT) (left panel) and plasma insulin levels (right panel) (*n*=8 per group). (I) Insulin tolerance test (ITT). Absolute blood glucose levels (left panel) and relative blood glucose levels (right panel) are shown. (J-K) Cold tolerance test. Mice were housed at room temperature (RT, 22°C) and then in a cold room (6°C) for 6h. (J) Body temperatures (*n*=6 per group). (K) qRT-PCR of *Ucp1* and *Elovl3* in BAT (*n*=4 per group). All quantitative data for mice are presented as means ± SEM. Statistical comparison between groups was performed using Student’s *t* -test (* p < 0.05 and *\*\** P < 0.01).

Blood glucose and insulin levels increased in the serum of A-KO mice, while leptin levels decreased. Free fatty acid, triglyceride and cholesterol levels were similar between A-KO and f/f mice (Figure 1G). Glucose tolerance test (GTT) and insulin tolerance test (ITT) showed that A-KO mice were insulin resistant (Figure 1H-I). We also observed reduced lipolysis and increased total energy expenditure upon stimulation with CL 316,243 (CL), a β3 adrenergic receptor agonist, in A-KO mice (Figure S2A-B). Deletion of *Med1* in adipocytes did not affect food intake and total activity (Figure S2C). A-KO mice maintained core body temperature when housed at room temperature (~22 °C), but their body temperature significantly dropped to ~31 °C after 3 h exposure to an ambient temperature of 6 °C (Figure 1J). Expression of thermogenesis genes *Ucp1* and *Elovl3* decreased in BAT of A-KO mice at room temperature and failed to induce after 3 h cold exposure, indicating thermogenesis defects in A-KO mice (Figure 1K). These data indicate that adipocyte-selective deletion of *Med1* impairs glucose homeostasis and energy expenditure in mice.

Under a high fat diet (HFD), A-KO mice gained significantly less body weight and fat mass compared to f/f mice, but more lean mass (Figure 2A-C and Figure S3A-B). *Med1* KO had little effect on food intake and energy expenditure (Figure 2D-E). After 8 wk of HFD, A-KO mice exhibited severe lipodystrophy phenotypes including dramatically reduced iWAT and eWAT mass, fatty liver, insulin resistance, hyperlipidemia and reduced serum leptin levels (Figure 2F-L and Figure S3C-E). These data indicate that mice with adipocyte-selective deletion of *Med1* are resistant to HFD-induced obesity.

**Figure 2.**
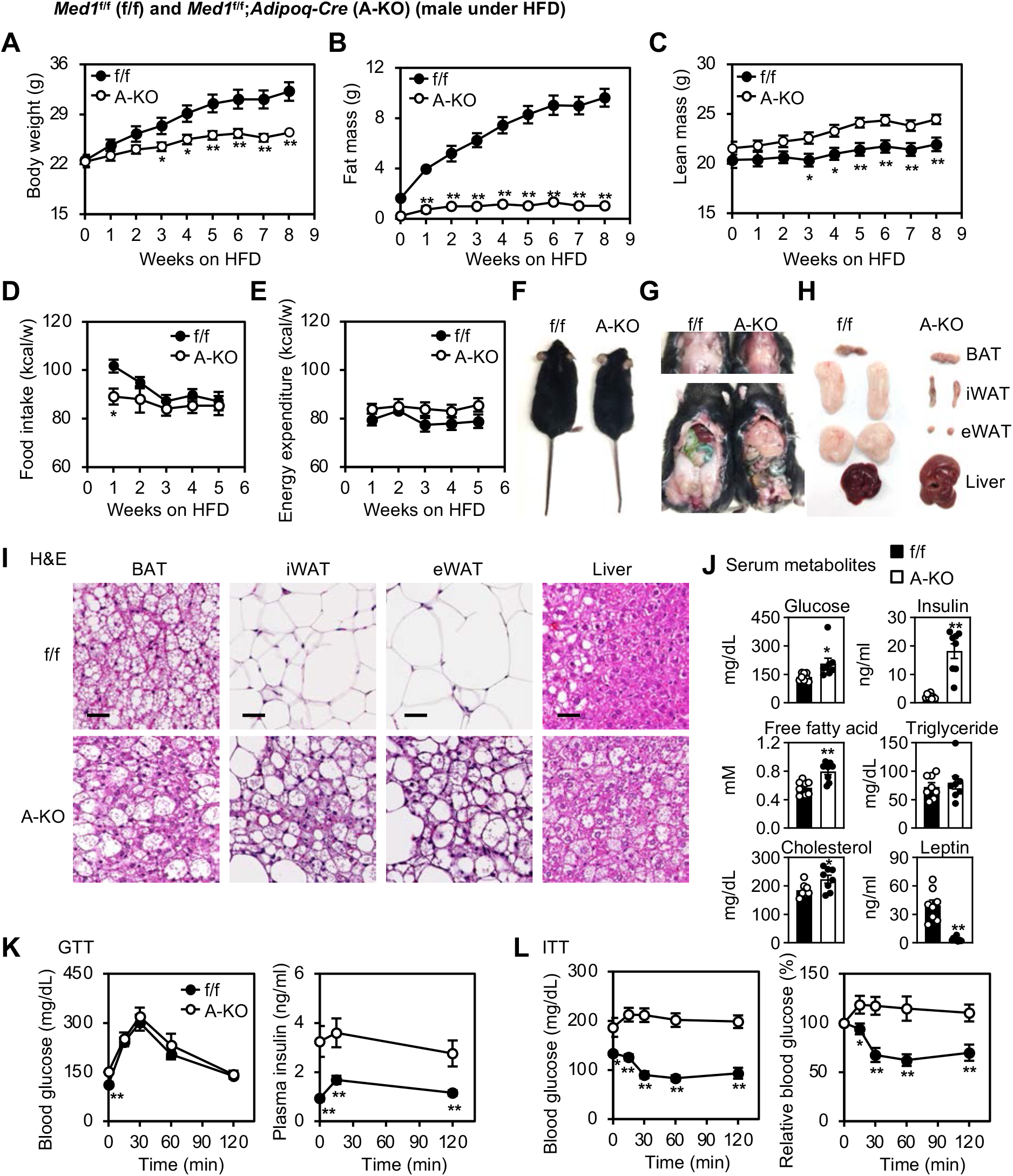
Mice with adipocyte-selective deletion of *Med1* are resistant to high fat diet-induced obesity. Male f/f and A-KO mice (*n*=8 per group) were fed with a high fat diet (HFD) from the 8th week of age. Data for female mice are shown in Figure S3. (A-C) Total body weight (A), fat mass (B) and lean mass (C) were measured by MRI during HFD feeding. (D-E) Food intake (D) and total energy expenditure (E). (F) Representative morphology of HFD-fed mice. (G) Representative pictures of the interscapular area (upper area) and abdominal area (lower panel) after 8wk of HFD feeding. (H) Representative pictures of each fat depot and liver. (I) H&E staining of each fat depot and liver. *Scale bar =* 100 μm. (J) Levels of serum metabolites. (K) GTT (left panel) and plasma insulin levels (right panel). (L) ITT. Absolute blood glucose levels (left panel) and relative blood glucose levels (right panel) are shown. Statistical comparison between groups was performed using Student’s *t* -test (* P < 0.05 and ** P < 0.01).

### MED1 is dispensable for embryonic development of adipose tissue

To find out whether MED1 is required for adipose tissue development, we used *Myf5-Cre* mice to delete *Med1* in precursor cells of BAT and skeletal muscle^26^. *Med1^f/f^;Myf5-Cre* (M-KO) embryos were obtained at expected ratio without showing morphological differences compared to f/f embryos (Figure 3A-B). M-KO embryos did not show observable differences in BAT mass compared to f/f embryos (Figure 3C-D). In addition, Myf5-Cre-mediated deletion of *Med1* had little effects on the expression of adipogenesis markers *Pparg, Cebpa* and *Fabp4* but significantly reduced brown adipocyte marker *Ucp1* expression (Figure 3E). Decreased *Ucp1* expression in *Med1* KO BAT is consistent with previous observations that MED1 is required for *Ucp1* expression in brown adipocytes in culture^27,28^. Although M-KO mice had largely intact BAT and muscle before birth, they became runts and showed severe growth retardation and reduced survival rates around 3-4 wk after birth, possibly due to muscle development defects (Figure 3A and Figure S4). These data suggest that MED1 is largely dispensable for embryonic development of adipose tissue.

**Figure 3.**
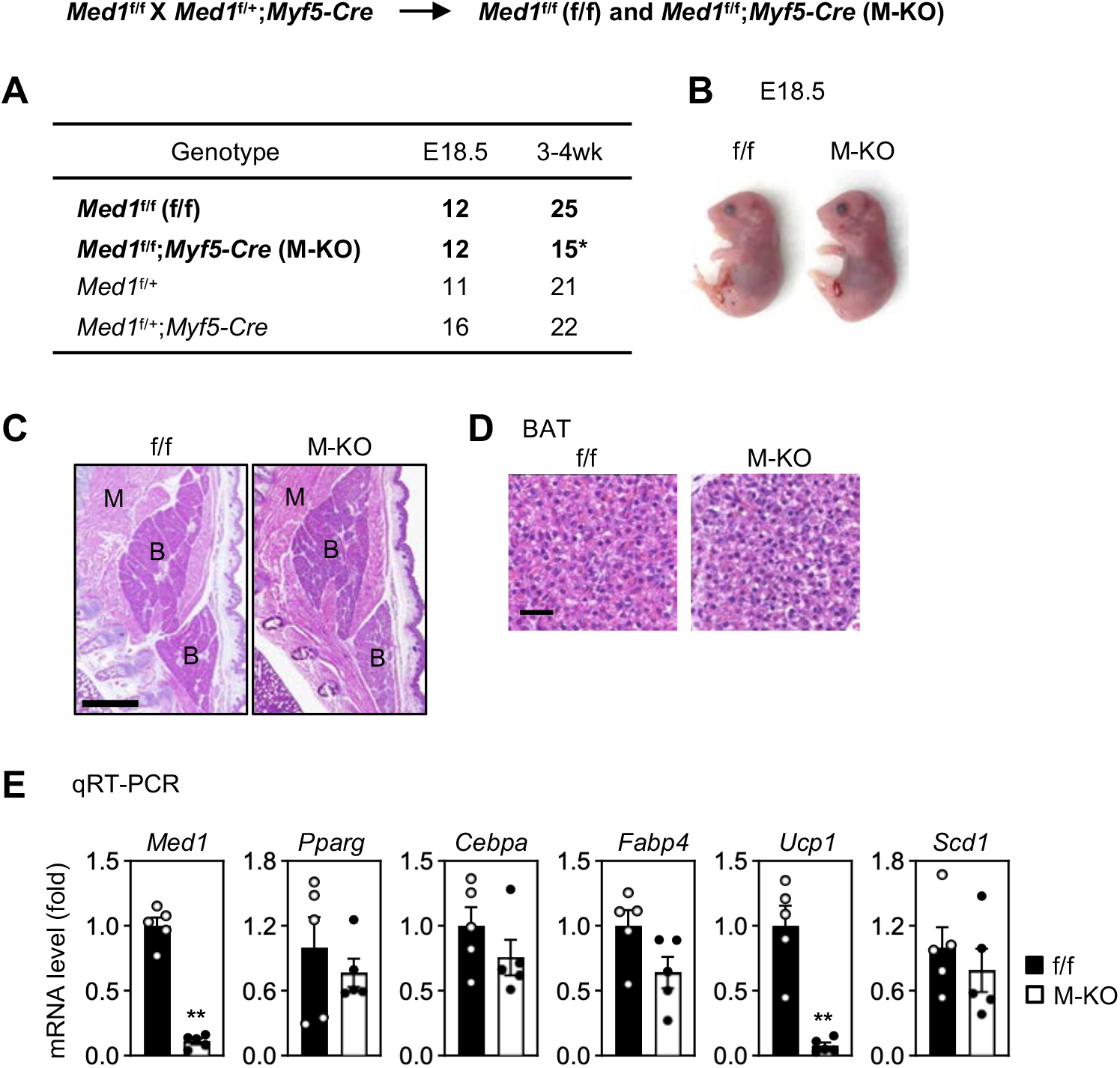
MED1 is dispensable for embryonic development of adipose tissue. (A-E) *Med1^f/f^;Myf5-Cre* (M-KO) mice show largely intact BAT before birth. *Med1^f/f^* (f/f) mice were crossed with *Myf5-Cre* to generate M-KO mice. (A) Genotypes of mice at E18.5 and 3-4wk. The expected ratio of the four genotypes is 1:1:1:1. Numbers of M-KO mice at 3-4wk were reduced as indicated by an asterisk. Data from 3-4wk M-KO mice are shown in Figure S4. (B) Representative morphology of E18.5 embryos. (C) H&E staining of E18.5 embryos. Sagittal sections of the cervical/thoracic area are shown. *Scale bar* = 80 μm. (D) H&E staining of BAT at E18.5. (E) qRT-PCR of *Med1, Pparg, Cebpa, Fabp4, Ucp1* and *Scd1* expression in BAT of E18.5 embryos (*n*=5 per group). Statistical comparison between groups was performed using Student’s *t* -test (* P < 0.05 and ** P < 0.01).

### MED1 is required for postnatal adipose expansion and lipogenesis gene expression

Since the requirement for MED1 to maintain adiposity was observed only after birth, we focused on postnatal adipose expansion in mice, which is stimulated by changes from breastfeeding to carbohydrate-rich diet (Figure 4A)^6^. We sought to profile cell type-specific transcriptomes in adiponectin-positive *(Adipoq^+^)* adipocytes during postnatal adipose expansion (Figure 4B-C). For this purpose, we first crossed *TRAP* (Translating Ribosome Affinity Purification) mice^29^ with *Adipoq-Cre* mice to label ribosomes in *Adipoq^+^* adipocytes. GFP-tagged ribosomes were isolated from interscapular BAT and iWAT of newborn and adult mice for RNA-Seq analyses (Figure 4D). Using 5-fold cut-off for differential gene expression, we identified genes that are induced (1.6%, 214/13,285) or reduced (1.8%, 240/13,285) in *Adipoq^+^* brown adipocytes from postnatal day 0.5 (P0.5) to 12 wk (Figure 4E). Induced genes were functionally associated with lipogenesis but not adipogenesis. Lipogenesis genes including *Acly, Acaca, Fasn* and *Scd1,* which encode four key fatty acid synthesis enzymes, were markedly induced in brown adipocytes from newborn to adult mice (Figure 4E-F). Similarly, we identified induced (4.0%, 552/13,799) or reduced (5.4%, 745/13,799) genes in *Adipoq^+^* white adipocytes from P2.5 to 12wk (Figure S5A). Four key lipogenesis enzyme genes *Acly*, *Acaca*, *Fasn* and *Scd1* were induced over 5-fold in white adipocytes from P2.5 to 12wk (Figure S5A-C). Western blot analyses confirmed that protein levels of SCD1 and FASN were highly induced during postnatal expansion of BAT and iWAT, but not those of adipogenesis markers and master regulators PPARγ and C/EBPα (Figure 4G). These results indicate that postnatal adipose expansion is associated with marked induction of lipogenesis enzymes in adipocytes from newborn to adult mice.

**Figure 4.**
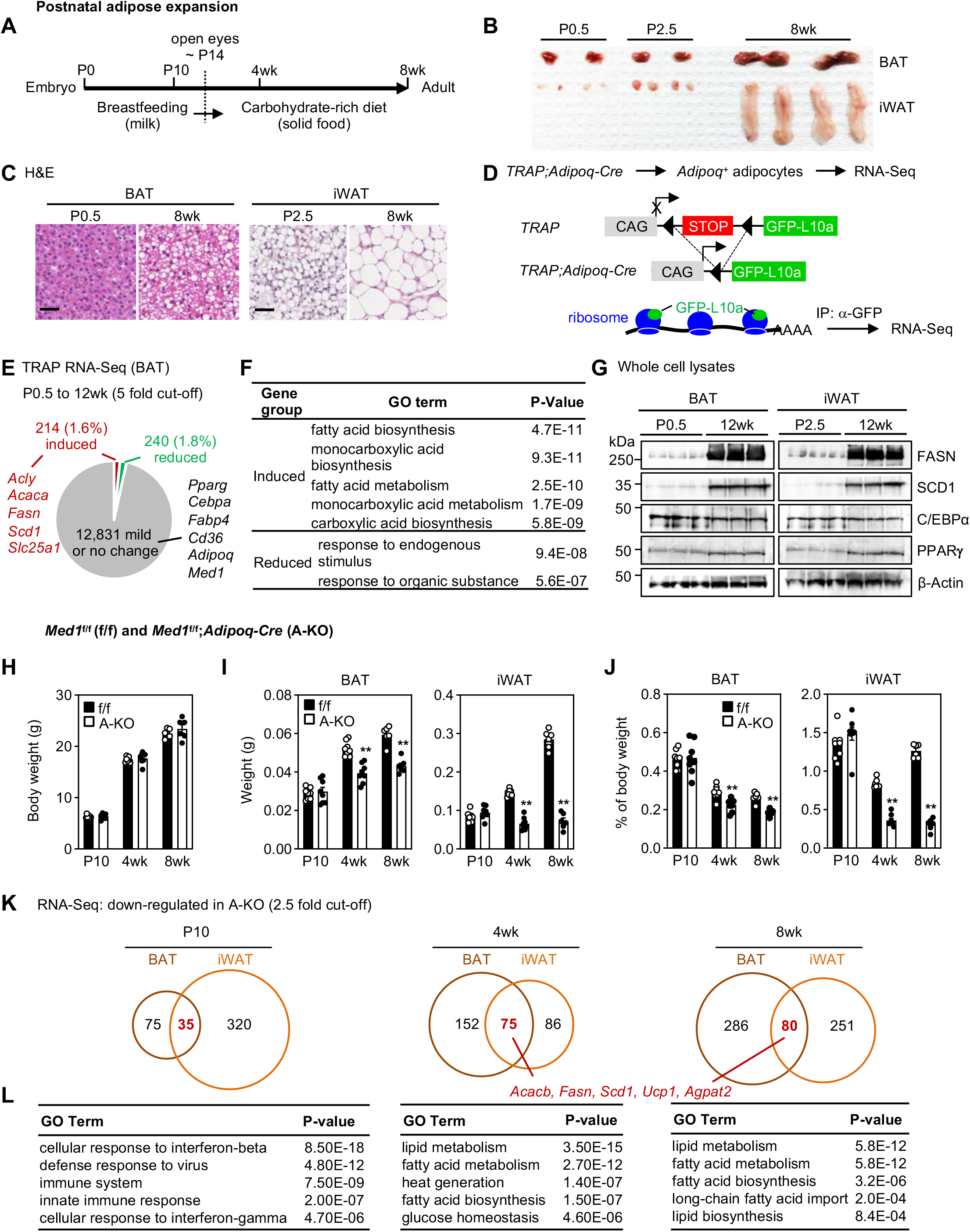
MED1 is required for postnatal adipose expansion and lipogenesis gene expression. (A-C) Increased adipose tissue size and lipid accumulation from newborn to adult mice. (A) Schematics of postnatal development stages from newborn to adult mice. (B) Representative pictures of BAT and iWAT of mice at postnatal day 0.5 (P0.5), P2.5 and 8wk. Adipose tissues from two mice are shown. (C) H&E staining of BAT and iWAT. *Scale bar* = 80 μm. (D-F) Lipogenesis gene expression is highly induced during postnatal expansion of BAT. iWAT data are shown in Figure S5. (D) Schematics of experimental design. *TRAP* mice were crossed with *Adipoq-Cre* to delete the STOP signal and generate mice expressing GFP-fused L10a, an integral component of the 60S ribosomal subunit, in *Adipoq^+^* adipocytes. GFP-L10a tagged ribosomes were immunoprecipitated with GFP antibody and mRNA was purified for RNA-Seq. (E) RNA-Seq analysis of *Adipoq^+^* brown adipocytes isolated from BAT of *TRAP;Adipoq-Cre* mice at P0.5 and 12wk. The cut-off for induced or reduced genes from P0.5 to 12wk is 5-fold. (F) Gene ontology (GO) analysis of gene groups defined in (E). (G) Western blot analysis of FASN, SCD1, C/EBPα and PPARγ using whole cell lysates from BAT (left panel) or iWAT (right panel). β-Actin was used as loading control. (H-L) MED1 is required for postnatal adipose expansion and lipogenesis gene expression during carbohydrate-rich diet feeding stages. (H-J) Total body weight (H), average tissue weight (I) and % of body weight (J) are presented (*n*=8 per group at P10 and 4 wk; *n*=6 at 8 wk). Statistical comparison between groups was performed using Student’s *t* -test (** P < 0.01). (K-L) BAT and iWAT were collected from f/f and A-KO mice at P10, 4wk and 8wk. Total RNA isolated from 3-5 mice per genotype was combined in equal amounts for RNA-Seq. (K) Numbers of down-regulated genes in adipose tissues of A-KO mice. The cut-off for differential expression is 2.5-fold. (L) GO analysis of down-regulated genes defined in (K) in both BAT and iWAT of A-KO mice are shown.

To investigate the role of MED1 in postnatal adipose expansion, we isolated fat depots from f/f and A-KO mice at breastfeeding stage (P10) and carbohydrate-rich diet feeding stages (4wk and 8wk). We did not observe any discernible differences in total body weight and interscapular WAT between f/f and A-KO mice at P10 (Figure 4H and Figure S6A). However, A-KO mice exhibited nearly complete loss of interscapular WAT as well as reduced BAT and iWAT mass at 4wk and 8wk (Figure 4I-J and Figure S6A). These results suggest that MED1 is required for adipose expansion during carbohydrate-rich diet feeding stages, but not during breastfeeding. RNA-Seq analysis showed that Adipoq-Cre-mediated deletion of *Med1* impaired the induction of lipogenesis genes including *Acly*, *Acaca*, *Acacb*, *Fasn*, and *Scd1* from P10 to 4wk and 8wk in both BAT and iWAT but did not affect the expression of adipogenesis marker *Pparg* (Figure 4K-L and Figure S6B). Together, these data indicate that MED1 is required for postnatal adipose expansion and lipogenesis gene expression when mice switch from breastfeeding to a carbohydrate-rich diet.

### MED1 is required for lipogenesis but not early adipogenesis in culture

To understand how MED1 regulates lipogenesis, we established inducible *Med1* KO *(Med1^f/f^;Cre-ER)* brown preadipocytes. Cells were treated with 4-hydroxytamoxifen (4OHT) to delete *Med1,* followed by the induction of adipogenesis. Consistent with A-KO mouse phenotype, *Med1* KO adipocytes showed more than 50% decrease in lipid content at day 7 (D7) of adipogenesis (Figure 5A). RNA-Seq analysis confirmed deletion of exons 8-10 of *Med1* gene (Figure S7A). MED1 is a subunit of general transcription coactivator complex Mediator. However, only about 1% of expressed genes were down-regulated in *Med1* KO adipocytes at D7 (Figure 5B). GO analysis showed that down-regulated genes were strongly functionally associated with fatty acid biosynthesis (Figure 5C). Consistent with data obtained from mouse adipose tissues, deletion of *Med1* led to decreases in expression of lipogenesis genes *Acly, Acacb, Fasn* and *Scd1* but not adipogenesis marker *Pparg* in culture (Figure S7B-C). Similar results were obtained with primary brown and white adipocytes differentiated in culture (Figure S8).

**Figure 5.**
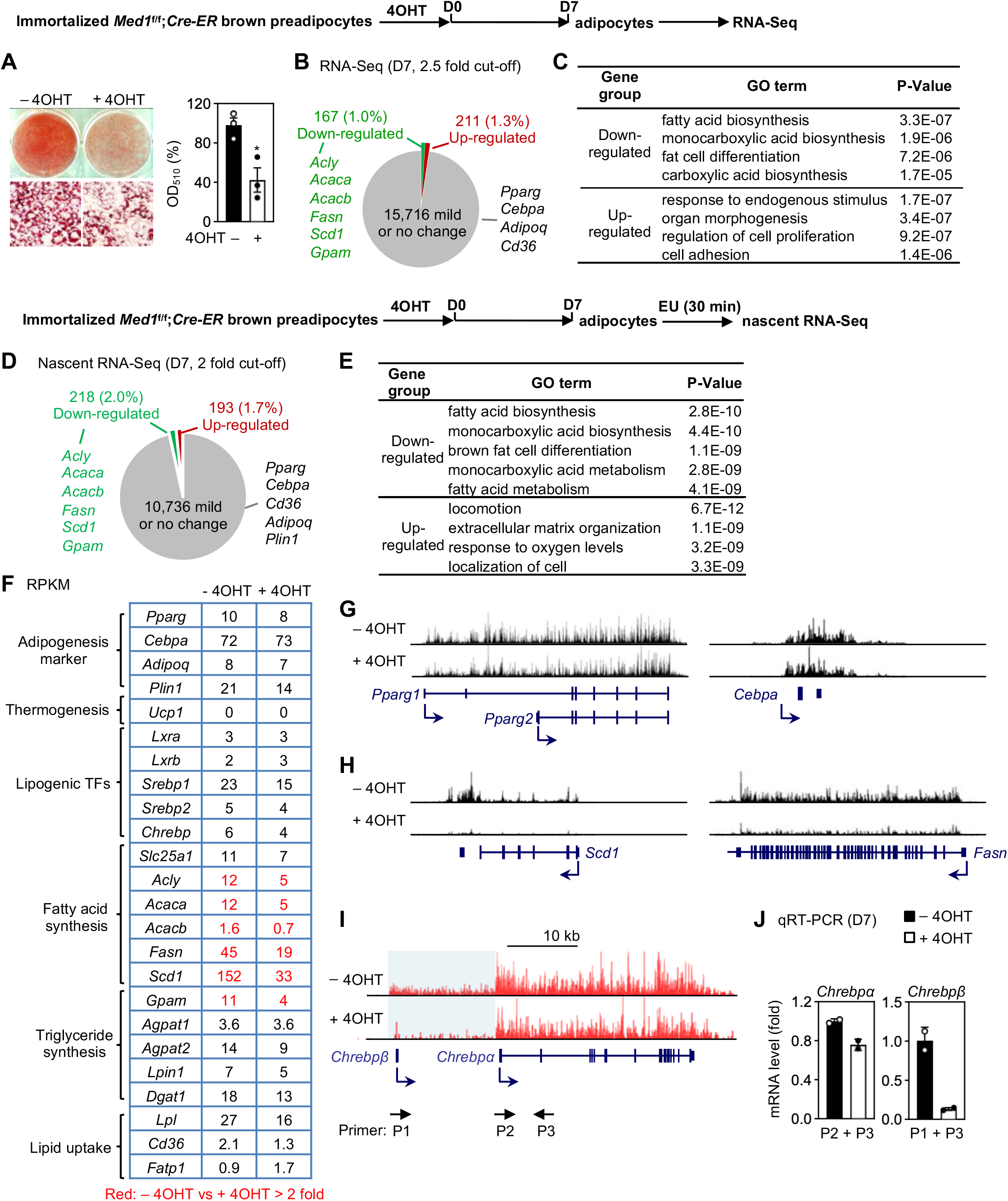
MED1 is required for lipogenesis but not early adipogenesis in culture. Immortalized *Med1^f/f^;Cre-ER* brown preadipocytes were treated with 4OHT to delete *Med1,* followed by adipogenesis assay, RNA-Seq or nascent RNA-Seq. (A-C) MED1 is required for lipid accumulation and lipogenesis gene expression in culture. (A) Oil Red O staining at day 7 (D7) of adipogenesis is shown (left panel). Lipid accumulation was quantified by extracting Oil Red O using isopropanol and reading absorbance at 510 nm (right panel). *n*=3 biological replicates. Data are presented as means ±SD. Statistical comparison between groups was performed using Student’s *t* -test (* P < 0.05). (B) RNA-Seq analysis at D7 of adipogenesis. The cut-off for differential expression is 2.5-fold. (C) GO analysis of gene groups defined in (B). (D-H) MED1 is required for transcription of lipogenesis genes. Cells were treated with Ethylene Uridine (EU) for 30min at D7 of adipogenesis for nascent RNA-Seq analysis. (D) Nascent RNA-Seq analysis at D7 of adipogenesis. The cut-off for differential expression is 2-fold. (E) GO analysis of gene groups defined in (D). (F) Expression levels of representative genes are shown in RPKM values. (G-H) Profiles of nascent RNA-Seq data are shown around adipogenesis genes *Pparg* and *Cebpa* (G) and lipogenesis genes *Scd1* and *Fasn* (H) loci. (I-J) Decreased *Chrebpβ* expression in *Med1* KO brown adipocytes. (I) Profiles of nascent RNA-Seq data around *Chrebp* locus. (J) qRT-PCR of *Chrebpα* and *Chrebpβ* expression at D7 of adipogenesis. All qPCR data in cells are presented as means ± SD. Two technical replicates from a single experiment were used.

To better assess the role of MED1 in regulating lipogenesis gene transcription, we performed nascent RNA-Seq analysis. Using a 2-fold cut-off, we found that only about 2% of expressed genes were down-regulated transcriptionally in *Med1* KO adipocytes at D7 (Figure 5D). Consistent with steady-state RNA-Seq data, nascent RNA-Seq analysis indicated that deletion of *Med1* reduces transcription levels of lipogenesis genes *Fasn* and *Scd1* but not adipogenesis markers *Pparg* and *Cebpa* (Figure 5E-H). Interestingly, while *Med1* KO did not affect the transcription of lipogenic TFs LXRα, LXRβ and SREBP1, it specifically reduced transcription of *Chrebpβ* isoform, which encodes a key lipogenic TF in adipocytes^10,13^ (Figure 5I-J), suggesting that ChREBP may play a role in MED1-dependent lipogenesis. Together, these results suggest that MED1 is required for lipogenesis but not early adipogenesis in culture.

### MED1 is required for Pol II binding on lipogenesis genes in adipocytes

To investigate whether MED1 affects chromatin accessibility and enhancer activation, we performed ATAC-Seq (<u>A</u>ssay for <u>T</u>ransposase <u>A</u>ccessible <u>C</u>hromatin with high-throughput sequencing) and ChIP-Seq analyses of active enhancer mark H3K27ac, and RNA Polymerase II (Pol II) at D7 of adipogenesis. On Med1^+^ promoters or enhancers in adipocytes, deletion of *Med1* did not affect chromatin accessibility or H3K27ac levels (Figure 6A-B). However, *Med1* KO led to decreased binding of serine-5 phosphorylated (S5P) initiating Pol II, while having limited effects on serine-2 phosphorylated (S2P) elongating Pol II (Figure 6A-B)^30^. GO analysis revealed that genes associated with MED1-dependent S5P-Pol II were strongly functionally associated with lipid metabolism including fatty acid biosynthesis (Figure 6C). Consistent with nascent RNA-Seq data, deletion of *Med1* clearly reduced both S5P-Pol II and S2P-Pol II binding on *Scd1, Fasn, Acaca* and *Gpam* loci but not the *Pparg* locus (Figure 6D and Figure S9). These data indicate that while MED1 is dispensable for chromatin opening and enhancer activation, it is required for Pol II binding on lipogenesis genes in adipocytes.

**Figure 6.**
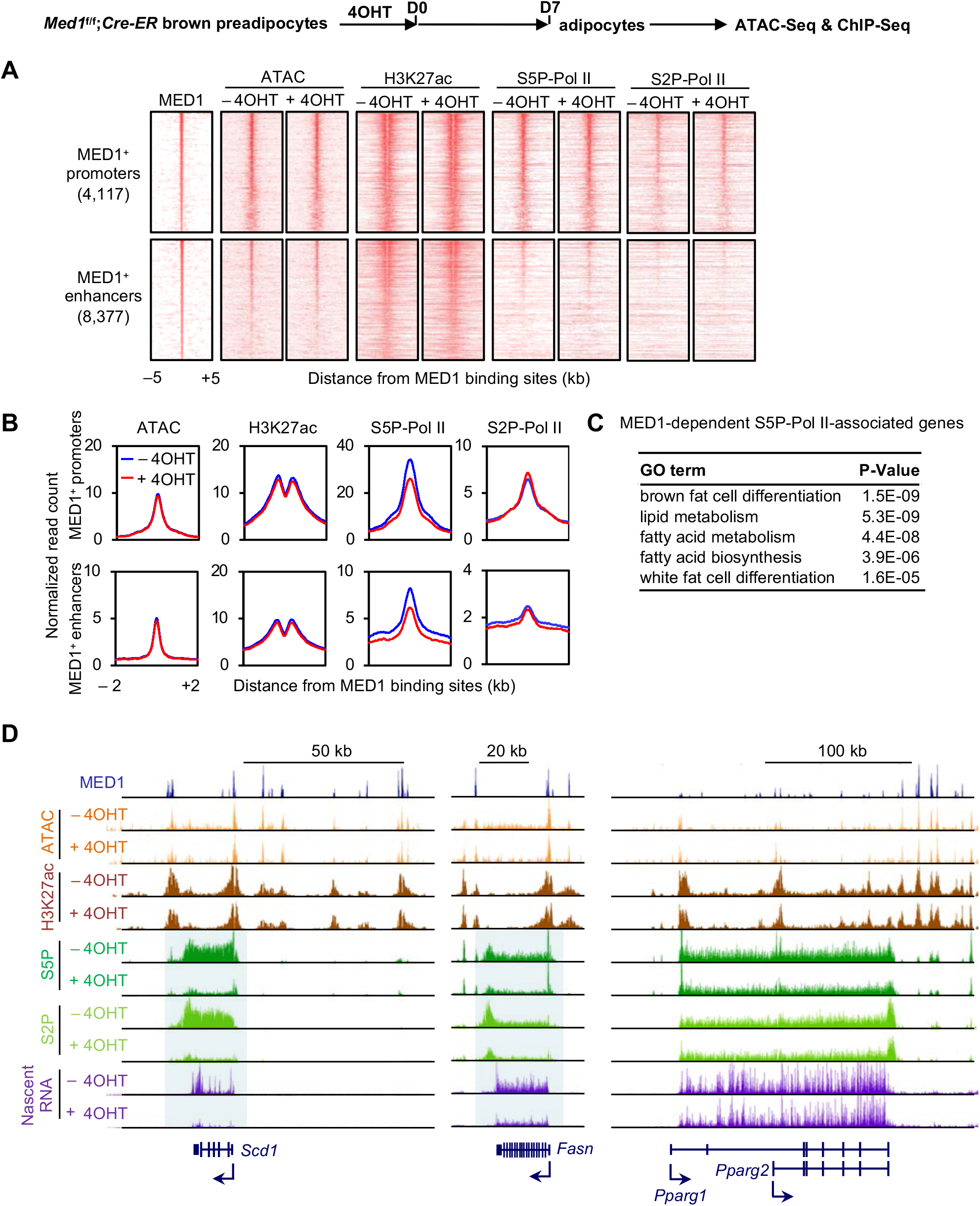
MED1 is required for Pol II binding on lipogenesis genes in adipocytes. Adipogenesis was done as in Figure 5. Cells were collected at D7 of adipogenesis for ATAC-Seq and ChIP-Seq of H3K27ac, S5P-Pol II and S2P-Pol II. (A-B) *Med1* KO does not affect chromatin accessibility or enhancer activation but reduces S5P-Pol II binding on MED1^+^ promoters and enhancers in adipocytes. Heat maps (A) and average profiles (B) were aligned around the center of MED1 binding sites on MED1^+^ promoters (4,117) and enhancers (8,377). Published MED1 ChIP-seq data was used (GSE74189)^44^. (C) GO analysis of genes associated with MED1-dependent S5P-Pol II. (D) MED1 is required for Pol II binding on lipogenesis genes. Profiles of ATAC-Seq, H3K27ac enrichment, S5P-Pol II, S2P-Pol II binding and nascent RNA-Seq data around *Scd1, Fasn* and *Pparg* gene loci are shown. Profiles around additional lipogenesis genes are shown in Figure S9.

### MED1 facilitates Mediator binding on ChREBP^+^ lipogenic enhancers in adipocytes

We hypothesized that MED1 regulates Pol II binding on lipogenesis genes by facilitating Mediator binding on lipogenic enhancers. To test this hypothesis, we first identified lipogenic enhancers that are bound by ChREBP. *Med1*^f/f^;*Cre-ER* brown preadipocytes were infected with retroviruses expressing triple T7 tagged constitutive active form of ChREBP (T7-CA-ChREBP). Cells were treated with 4OHT to delete *Med1* (Figure 7A), followed by the induction of adipogenesis. We found that ectopic ChREBP failed to rescue the reduced lipogenesis in *Med1* KO adipocytes (Figure 7B-C). Next, we performed ChIP-Seq using antibodies against T7 and MED12, which is a representative Mediator subunit^31^, at D7 of adipogenesis. Motif analysis of the T7-CA-ChREBP binding regions identified the ChREBP motif as a top motif (Figure 7D). Among the 2,627 ChREBP^+^ active enhancers, 934 showed reduced MED12 binding in *Med1* KO adipocytes (Figure 7E-F). Notably, fatty acid metabolism was a top GO term associated with genes that showed MED1-dependent MED12 binding (Figure 7G). Moreover, ChREBP-associated genes that show MED1-dependent MED12 binding exhibited reduced expression in *Med1* KO cells (Figure 7H), indicating that MED1 facilitates MED12 binding on ChREBP^+^ lipogenic enhancers to regulate lipogenesis gene expression in adipocytes. Indeed, deletion of *Med1* clearly reduced MED12 binding to lipogenesis genes *Scd1, Fasn, Acaca* and *Acly* loci but not the *Pparg* locus in adipocytes (Figure 7I and Figure S10). Together with previous data, these results suggest that MED1 is a lipogenesis coactivator required for Mediator binding on ChREBP^+^ lipogenic enhancers in adipocytes (Figure 7J).

**Figure 7.**
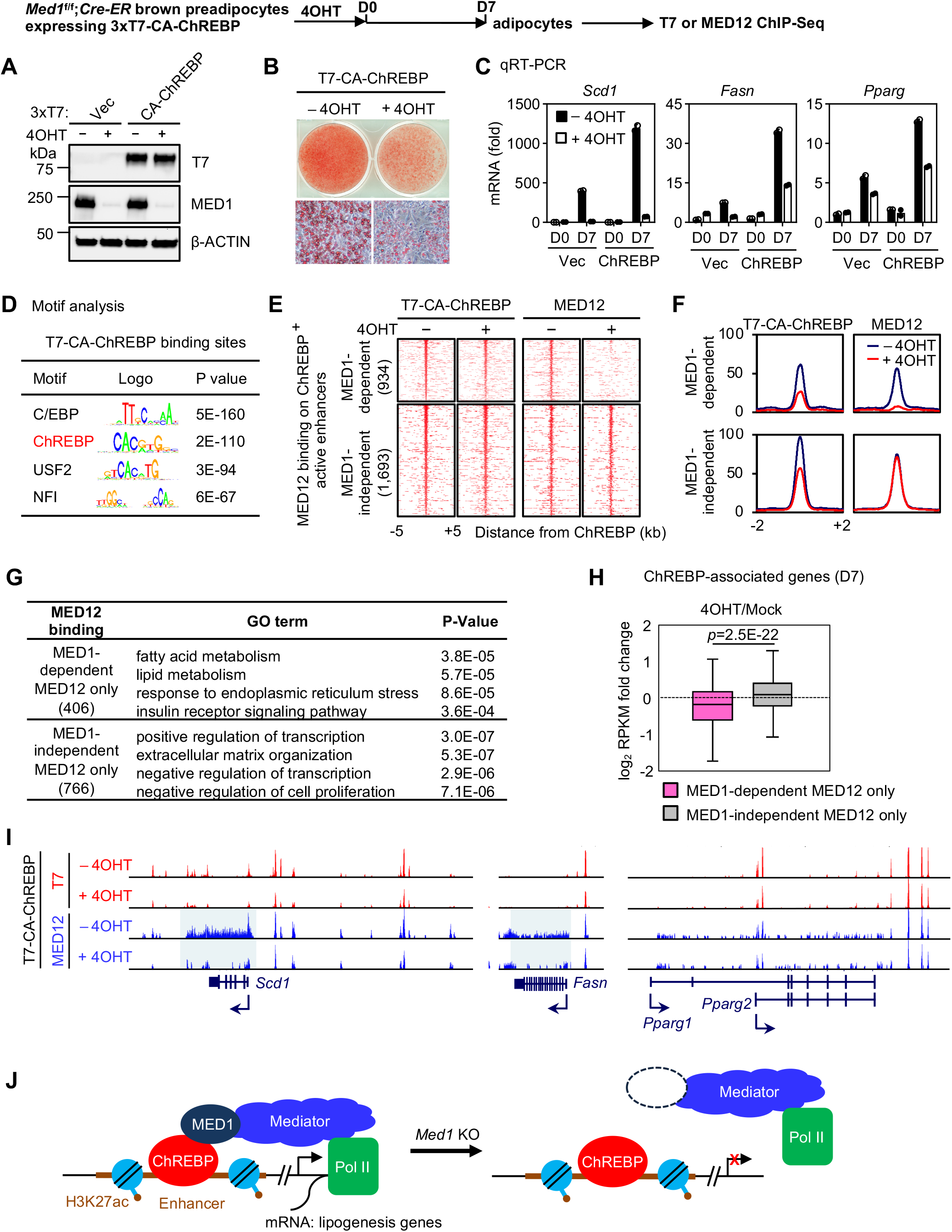
MED1 facilitates Mediator binding on ChREBP^+^ lipogenic enhancers in adipocytes. Immortalized *Med1^f/f^;Cre-ER* brown preadipocytes were infected with retroviruses expressing 3xT7-CA (constitutive active)-ChREBP or 3xT7 vector alone. After hygromycin selection, cells were treated with 4OHT to delete *Med1,* followed by adipogenesis. Cells were collected at D7 for T7 and MED12 ChIP-Seq analysis. (A-C) Ectopic CA-ChREBP fails to rescue reduced lipogenesis in *Med1* KO. (A) Ectopic expression of CA-ChREBP in *Med1* KO preadipocytes. Nuclear extracts were analyzed by Western blot using respective antibodies. (B) Oil Red O staining at D7 of adipogenesis. (C) qRT-PCR of *Scd1, Fasn* and *Pparg* expression at D7 of adipogenesis. Two biological replicates were used. (D) Motif analysis of top 3,000 T7-CA-ChREBP binding sites. (E-F) MED1 is required for MED12 binding on a subset of ChREBP^+^ active enhancers. Heat maps (E) and average profiles (F) around the center of T7-CA-ChREBP^+^ active enhancers are shown. (G) GO analysis of genes that are associated with either MED1-dependent or MED1-independent MED12 binding on ChREBP^+^ active enhancers. (H) MED1 facilitates MED12 binding to promote ChREBP-associated gene expression. Fold changes of RPKM values between control (Mock) and *Med1* KO (4OHT) cells are shown in box plots. Statistical significance levels are indicated (Wilcoxon signed rank test, two-sided). (I) MED1 is required for MED12 binding on lipogenesis genes. Profiles of T7-CA-ChREBP and MED12 binding on *Scd1, Fasn* and *Pparg* gene loci are shown. Profiles around additional lipogenesis genes are shown in Figure S10. (J) Proposed model depicting that MED1 is required for Mediator binding on ChREBP^+^ lipogenic enhancers in adipocytes.

## DISCUSSION

By crossing *Med1^f/f^* with *Adipoq-Cre* or *Myf5-Cre* mice, we selectively deleted *Med1* in adipocytes or precursor cells *in vivo.* We found surprisingly that MED1 is required for postnatal adipose expansion but is largely dispensable for the development of adipose tissues. Consistently, MED1 is required for lipid accumulation but not early differentiation during adipogenesis in culture. Transcriptome analysis in mice and in culture indicate that only 2-3% of expressed genes are down-regulated by *Med1* KO in adipocytes. Down-regulated genes include key lipogenesis enzymes such as SCD1 and FASN, but not early adipogenesis markers such as PPARγ. Nascent RNA-Seq in adipocytes reveals that *Med1* KO specifically reduces transcription of ChREBPβ isoform but not other lipogenic TFs. We further demonstrate that MED1 regulates Pol II binding on lipogenesis genes by promoting Mediator binding on ChREBP^+^ lipogenic enhancers in adipocytes. Together, our findings suggest that the MED1 subunit of the Mediator coactivator complex regulates postnatal adipose expansion by promoting lipogenesis gene expression.

By crossing *Med1^f/f^* with *Myf5-Cre* mice, we observed that *Med1* KO had little effects on the size of embryonic BAT and the expression of adipogenesis markers such as PPARγ and C/EBPα. A-KO mice showed similar BAT and iWAT tissue masses and adipogenesis marker expression as the control mice at P10. In culture, deletion of *Med1* does not affect PPARγ or C/EBPα expression during differentiation of white and brown adipocytes. Together, these results indicate that MED1 is largely dispensable for the general development of adipose tissues. The requirement of MED1 for *Ucp1* induction in BAT and iWAT is consistent with previous findings in cell culture that PRDM16 physically binds to and recruits MED1 to active enhancers of BAT-selective genes and that both PRDM16 and MED1 are required for *Ucp1* induction in brown adipocytes^27,28^. We also show that MED1 is largely dispensable for embryonic development of muscle. However, M-KO mice show severe growth retardation and mortality around weaning age, suggesting that MED1 may play a role in muscle development and function after birth. These observations suggest that MED1 plays functional roles in late stages of adipogenesis and myogenesis rather than early stages of cell differentiation and tissue development.

Using adipocyte-specific transcriptome analysis, we show that postnatal adipose expansion is associated with marked induction of fatty acid synthesis enzymes (Figure 4D-G and S5). Transcriptome analyses at P10, 4wk and 8wk also reveal marked induction of fatty acid synthesis enzymes as well as lipogenic TF *Chrebp* in BAT and iWAT when mice switch from maternal milk to a carbohydrate-based diet (Figure S6B). These results explain previous observations that lipogenic enzyme activity and fatty acid synthesis in adipose tissues increase markedly after mice switch from breastfeeding to a carbohydrate-based diet^6^. A-KO mice exhibit increasing lipodystrophy after weaning but not during the breastfeeding stage. A-KO mice also show severely impaired induction of fatty acid synthesis genes in BAT and iWAT when switching from high-fat maternal milk to a carbohydrate-based diet. These data suggest that MED1 is required for carbohydrate-rich diet-induced postnatal adipose expansion and that lipodystrophy in A-KO mice is at least in part due to impaired induction of fatty acid synthesis genes. Consistently, *Med1* KO reduces lipid accumulation and impairs the induction of fatty acid synthesis genes in adipocytes in culture.

Impaired induction of triglyceride synthesis and lipid uptake genes could also contribute to lipodystrophy in A-KO mice. AGPAT2 is a critical enzyme for triglyceride synthesis and is highly expressed in adipose tissues. Mutations of *AGPAT2* lead to congenital generalized lipodystrophy in humans^32^. Whole-body *Agpat2* KO mice show a severe lipodystrophy phenotype similar to A-KO mice^33^. A-KO mice show impaired induction of *Agpat2* in BAT and iWAT from P10 to 4wk (Figure S6B). *Med1* KO reduces *Agpat2* expression in primary white and brown adipocytes in culture (Figure S8G). These results suggest that the impaired induction of *Agpat2* likely contributes to the lipodystrophy in A-KO mice. We also observed decreased expression of lipid uptake genes *Fatp1* and *Fatp2* in adipose tissues of A-KO mice (Figure S6B). We cannot exclude the possibility that decreased expression of *Fatp1/2* contributes to lipodystrophy in A-KO mice.

In *Med1* KO adipocytes, only 1-2% of the ~16,000 expressed genes are down-regulated, indicating that MED1 is not generally required for transcription. Down-regulated genes are preferentially associated with lipogenesis in particular fatty acid synthesis, such as *Acaca*, *Elovl6, Fasn, Scd1* and *Chrebpβ.* ChREBP is a major transcriptional regulator of fatty acid synthesis in adipocytes. Adipoq-Cre-mediated KO of *Chrebp* reduces the expression of lipogenesis genes including *Acaca*, *Elovl6, Fasn* and *Scd1* in iWAT, although these mice show a milder lipodystrophy phenotype compared to A-KO mice^13^. This could be due to the difference in the genetic backgrounds between *Chrebp* KO and A-KO mice and the compensatory increase in *Srebp1* expression by *Chrebp* KO. SREBP1 is another critical transcriptional regulator for fatty acid synthesis in adipocytes. Overexpression of SREBP1a in adipose tissue increases the expression of fatty acid synthesis genes, leading to increased lipid accumulation in adipocytes^15^.

Previous studies have implicated other Mediator subunits in SREBP1-dependent lipogenesis gene expression. GST-fused SREBP1a activation domain can pull down the Mediator complex from cell nuclear extracts. SREBP1a directly interacts with MED15 (also called ARC105) *in vitro.* Depletion of MED15 down-regulates SREBP1a-dependent *FASN* expression in human cells^34^. CDK8 and Cyclin C, which associate with the transcriptionally inactive Mediator complex, have been reported as negative regulators of de novo lipogenesis in Drosophila and mice^35^. CDK8 and Cyclin C are dissociated from the Mediator complex upon re-feeding with a carbohydrate-rich diet after fasting^36^. Our finding that MED1 is a lipogenesis coactivator in adipocytes is consistent with these previous reports. Detailed molecular mechanisms by which Mediator regulates lipogenesis through its subunits including MED1, MED15, CDK8 and Cyclin C remain to be fully understood.

## MATERIALS AND METHODS

### Plasmids, antibodies and chemicals

Triple T7 (3xT7) double StrepII tag was N-terminally subcloned into retroviral vector pMSCVhyg to generate pMSCVhyg-3xT7 using pTAG1-hygroTK-N-term 3xT7 double StrepII tag (gift from Rob Klose)^37^ as a template. Constitutive active form of mouse ChREBP was subcloned into pMSCVhyg-3xT7 to generate pMSCVhyg-3xT7-CA-ChREBP using full length clone as templates (Addgene #39235). All plasmids were confirmed by DNA sequencing. Anti-MED1 (A300-793A) and anti-MED12 (A300-774A) were from Bethyl Laboratories. Anti-FASN (#3810), anti-SCD1(#2794) and anti-T7 (#132469) were from Cell Signaling Technology. Anti-C/EBPα (sc-61X) and anti-PPARγ (sc-7196X) were from Santa Cruz Biotechnology. Anti-H3K27ac (ab4729), anti-GFP (ab290), anti-S5P Pol II (ab5131) and anti-S2P Pol II (ab5095) were from Abcam. Anti-β-Actin (A1978) and (Z)-4-hydroxytamoxifen (4OHT) (H7904) were from Sigma.

### Generation of mouse strains

*Med1*^f/f^ mice^24^ were crossed with *Myf5-Cre* (Jackson no. 007893), *Adipoq-Cre* (Jackson no. 028020), or *Cre-ER* (Jackson no. 008463) to generate *Med1^f/f^;Myf5-Cre, Med1^f/f^;Adipoq-Cre,* or *Med1^f/f^;Cre-ER.* For genotyping of *Med1* alleles, PCR was performed using the following primers: 5’-TCTCCCCGGCTAATATTCATA-3’ and 5’-AAGGAACAAGCCAGCAAGC-3’. PCR amplified 839bp from the wild-type and 575bp from the floxed allele. *Rosa2^fssTRAP^ (TRAP)* mice (Jackson no. 022367)^29^ were crossed with *Adipoq-Cre* (Jackson no. 028020) to generate *TRAP;Adipoq-Cre* mice. For genotyping of *TRAP* alleles, PCR was performed using following primers: Common Forward, 5’-AAGGGAGCTGCAGTGGAGTA-3’; Wild-type Reverse, 5’-CCGAAAATCTGTGGGAAGTC-3’; Mutant Reverse, 5’-CGGGCCATTTACCGTAAGTTAT-3’. PCR amplified 197bp from the wild-type and 284bp from the *TRAP* allele. All mouse experiments were performed in accordance with the NIH Guide for the Care and Use of Laboratory Animals and approved by the Animal Care and Use Committee of NIDDK, NIH.

### Translating ribosome affinity purification

Brown and inguinal white adipose tissues from newborn (P0.5 for BAT or P2.5 for WAT) and adult (12wk) *TRAP;Adipoq-Cre* mice were minced in homogenization buffer (50 mM Tris [pH7.5], 12 mM MgCl2, 100 mM KCl, 1 % NP-40; 100 mg/ml cycloheximide, 1 mg/ml sodium heparin, 2 mM DTT, 0.2 units/ml RNasin, protease inhibitors) and homogenized by Dounce (Type B, 2 ml). After incubation on ice for 10 min, lysates were centrifuged at 13,000 × g for 10 min at 4 °C and the supernatant was transferred to new tubes after removing the top lipid layer. The supernatant was incubated with anti-GFP antibody (5 μg/ml) prebound to Dynabeads Protein A (Invitrogen) for 2h with end-over-end rotation to capture GFP-fused L10a subunit of the 60S ribosome. Beads were collected on a magnetic rack and washed twice with low-salt wash buffer (50 mM Tris [pH7.5], 12 mM MgCl2, 100 mM KCl, 1 % NP-40, 100 mg/ml cycloheximide, 2 mM DTT) and then three times with high-salt wash buffer (50 mM Tris [pH 7.5], 12 mM MgCl2, 300 mM KCl, 1 % NP-40, 100 mg/ml cycloheximide, 2 mM DTT). Immunoprecipitated ribosomes were immediately placed in the RLT buffer to extract RNA using the Qiagen Micro RNeasy kit (Qiagen) following the manufacturer’s protocol.

### Metabolic studies

For determination of serum metabolites, blood was collected from mice fed with standard laboratory mouse chow (7022 NIH-07, Harlan; 15% calories from fat, 56% calories from carbohydrate, 29% calories from protein) or high fat diet (D12492, Research Diets; 60% calories from fat, 20% calories from carbohydrate, 20% calories from protein). Plasma was obtained by centrifuging blood samples at 4 °C for 10 min at 12,000 × g. Plasma insulin and leptin concentrations were measured by an ELISA kit from CrystalChem and R&D Systems, respectively. Plasma free fatty acid, triglyceride and cholesterol concentrations were measured with reagents from Roche Diagnostics GmbH, Pointe Scientific Inc and Thermo Scientific, respectively. For GTTs, mice were fasted overnight for 12 h. For ITTs, mice were fasted for 4h. Mice received glucose (1g/kg i.p.; intraperitoneally), or human insulin (0.75 U/kg i.p., Humulin, Eli Lilly), respectively, and blood was collected from the tail vein at specific time points. For both tests, blood glucose levels were determined using a portable glucometer (Contour Glucometer, Bayer). For in vivo lipolysis, mice were injected i.p. with saline or CL316,243 (Sigma, 0.1 mg/kg) and blood was collected 20 min after the injection. Plasma concentrations of free fatty acid and glycerol were analyzed with reagents from Roche and Sigma, respectively. Food intake, O2 consumption, CO2 production were measured at 22°C over 24h period in a Comprehensive Lab Animal Monitor System (CLAMS) system (Columbus Instruments Inc.; 2.5L chambers with plastic floors, using 0.6 L/min flow rate, one mouse per chamber) after 48h adaptation period^38^. For the cold tolerance test, mice were individually housed at room temperature (RT, 22°C) and then in a cold room (6°C) for 6 h (n=6 per group). Core body temperature was measured using a rectal thermometer (TH-5, Braintree Scientific) before and hourly after cold exposure. Body composition was measured with the EchoMRI 3-in-1 analyzer (Echo Medical Systems).

### Primary preadipocytes culture, immortalization and adipogenesis

Primary brown preadipocytes were isolated from interscapular BAT of newborn *Med1^f/f^;Cre-ER* pups, and immortalized by infecting retroviruses expressing SV40T. Immortalized cells were further infected with retroviruses expressing 3xT7-CA-ChREBP or 3xT7 vector. Primary white adipocytes were from inguinal WAT of *Med1^f/f^;Adipoq-Cre* or *Med1^f/f^* adult mice. For adipogenesis assay, preadipocytes were plated in growth medium (DMEM plus 10% FBS) 3-4 days before the induction and were induced with 0.02 μM insulin, 1 nM T3, 0.5 mM IBMX, 2 μgml^−1^ DEX and 0.125 mM indomethacin for 2 days. For adipogenesis of primary white preadipocytes, 1 μM rosiglitazone was included throughout the differentiation^39,40^.

### Western blot and qRT-PCR

Western blot of nuclear extracts or whole tissue lysates was done as described^41^. Total RNA was extracted using TRIzol (Invitrogen) and reverse transcribed using ProtoScript II first strand cDNA synthesis kit (NEB), following the manufacturers’ protocols. qRT-PCR of *Med1* exon 8 was performed using SYBR green primers: forward 5’-CCTGTTTGATGGGATGTCCA-3’ and reverse, 5’-GCAGAGATATGCAGATTGCC-3’. *Chrepa* and *Chrebpβ* qRT-PCR primers were described previously^10^. SYBR green primers for other genes were described previously^39,42^.

### RNA-Seq library preparation

Total RNA (1 μg) or TRAP-isolated RNA (200 ng) was subjected to the NEBNext Poly(A) mRNA Magnetic Isolation Module (NEB) to isolate mRNA and proceeded directly to double stranded cDNA synthesis. Library construction was done using the NEBNext Ultra™ II RNA Library Prep Kit for Illumina (NEB) following the manufacturer’s protocol.

### Nascent RNA-Seq library preparation

10^6^ cells were labeled with 0.5 mM ethylene uridine for 30 min at 37 °C. After RNA extraction, 1 μg total RNA was depleted of rRNA using the NEBNext rRNA Depletion kit (NEB). To capture nascent RNA, the sample was biotinylated by the Click-iT Nascent RNA Capture Kit (ThermoFisher, C10365) according to the manufacturer’s instructions. Double-stranded cDNAs were synthesized using SuperScript Double-Stranded cDNA Synthesis Kit (Invitrogen). Library construction was done using NEBNext Ultra™ II DNA Library Prep Kit for Illumina (NEB) according to the manufacturer’s instructions. Sequencing libraries were analyzed with Qubit and pooled and sequenced on a HiSeq3000.

### ChIP-Seq and ATAC-Seq library preparation

ChIP-Seq was performed as described in detail previously^38,43–45^. ChIP-Seq of T7 and MED12 were done in the presence of *Drosophila* spike-in chromatin and antibody following the manufacturers’ protocol (Active Motif). ChIP-Seq library construction was done using NEBNext Ultra™ II DNA Library Prep Kit for Illumina (NEB) following the manufacturer’s protocol. For ATAC-Seq, 50,000 cells were washed with PBS, and collected in a cold lysis buffer (10 mM Tris-HCl, pH 7.4, 10 mM NaCl, 3 mM MgCl_2_ and 0.1% IGEPAL CA-630). After lysis, the nuclear pellet was resuspended in the transposase reaction mix (Illumina, Nextera Tn5 Transposase kit) and incubated for 30 min at 37°C. Immediately following transposition, the sample was purified by a Qiagen MinElute kit (Qiagen). Eluted DNA was amplified with PCR using Nextera i7- and i5-index primers (Illumina) and purified with AMPure XP magnetic beads (Beckman Coulter). Sequencing libraries were analyzed with Qubit and pooled and sequenced on a HiSeq2500 or HiSeq3000.

### Computational analysis

#### RNA-Seq data analysis

Raw sequencing data were aligned to the mouse mm9 genome using STAR software^46^. Reads on exonic regions or gene bodies (nascent RNA-Seq) were collected to calculate reads per kilobase per million (RPKM) as a measure of gene expression level. Only genes with RPKM > 1 were considered expressed. Gene ontology (GO) analysis of differentially expressed genes was carried out using DAVID bioinformatics resources (https://david.ncifcrf.gov).

#### ChIP-Seq peak calling and GO analysis of genomic regions

Raw sequencing data were aligned to the mouse mm9 genome using bowtie2. To identify ChIP-enriched regions, we used the ‘SICER’ method^47^. For H3K27ac enrichment, the window size of 200 bp and the estimated false discovery rate (FDR) threshold of 10^−10^ were used. For ChIP-Seq of S5P-Pol II, S2P-Pol II, T7, and MED12, the window size of 50 bp and the FDR threshold of 10^−3^ were used. For PPARγ peaks, the window size of 50 bp and the FDR threshold of 10^−10^ were used. Previously published MED1 ChIP-Seq data was used (GSE74189). GO analysis of genomic regions was done using GREAT (http://great.stanford.edu/public/html/).

#### Motif analysis

For motif analysis of 3xT7-CA-ChREBP binding sites, we used the SeqPos motif tool in Galaxy Cistrome (http://cistrome.org/ap/root) with default parameters. We selected the top 3,000 binding regions to screen enriched TF motifs based on the FDR value provided by SICER.

#### Normalization of ChIP-Seq data

For ChIP-Seq spike-in normalization, sequences were aligned to the *Drosophila* genome dm6. Normalization factors were determined by counting *Drosophila* tags and applying to mouse tags to generate normalized heat maps and profiles (Figure 7).

#### Heat maps and box plots

Heat map matrices were generated using in-house scripts with 50 bp resolution and visualized in R using gplots package. The ratio of RPKM values in *Med1* KO and control cells in base 2 logarithm was plotted using box plot, with outliers not shown (Figure 7). Wilcoxon signed rank test (two-sided) was used to determine statistical differences.

#### ATAC-Seq data analysis

Raw sequencing reads were processed using Kundaje lab’s ataqc pipelines (https://github.com/kundajelab/atac_dnase_pipelines). For downstream analysis, we used filtered reads that remained after removing unmapped reads, duplicates and mitochondrial reads.

## DATA AVAILABILITY

RNA-Seq, ATAC-Seq, ChIP-Seq datasets generated in the paper have been deposited in NCBI Gene Expression Omnibus under accession number GSE160605.

## SUPPLEMENTARY DATA

Supplementary Data are available online.

## FUNDING

This work was supported by the Intramural Research Program of NIDDK, NIH to KG.

## Conflict of interest statement

None declared.

## ACKNOWLEDGEMENTS

We thank Janardan K. Reddy for kindly providing *Med1-flox* mice, Aaron Broun and David Wu for technical assistance in genotyping, Rob Klose for 3xT7 tag plasmid, NIDDK Genomics Core and NHLBI DNA Sequencing and Genomics Core for NGS sequencing, Danyang Wang and Susanna Maisto for proofreading the manuscript.

## AUTHOR CONTRIBUTIONS

Conceptualization, Y.J. and K.G.; Methodology, Y.J., Y.-K.P., J.-E.L. and O.G.; Investigation, Y.J., Y.-K.P., J.-E.L., O.G., N.T. and K.G.; Software, Formal Analysis, and Data Curation, J.-E.L., Y.J. and Y.-K.P.; Writing – Original Draft, Y.J., J.-E.L. and K.G.; Writing – Review & Editing, Y.J., J.-E.L., Y.-K.P., N.T. and K.G.; Project Administration and Funding Acquisition, K.G.

---

**Figure S1.**
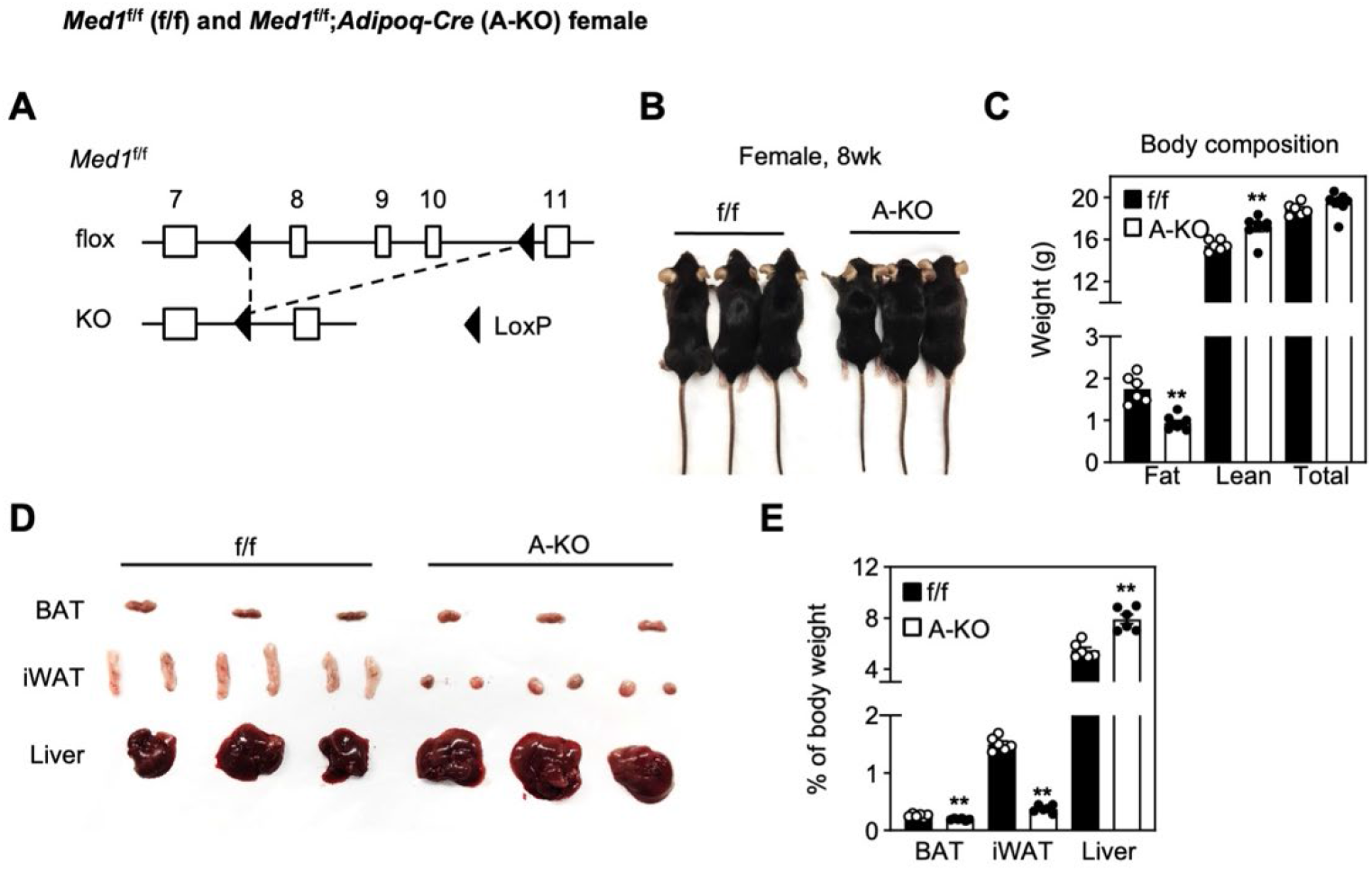
Related to Figure 1. *Med1*^f/f^;*Adipoq-Cre* female mice show severe lipodystrophy. 8 weeks (8wk) female *Med1*^f/f^ (f/f) and *Med1^f/f^;Adipoq-Cre* (A-KO) mice were fed with a regular diet (*n*=9 per group). (A) Schematics of conditional KO (flox) allele and KO allele for f/f mice targeting the exon 8-10 of *Med1^1^.* (B) Representative morphology of adult female mice. (C) Body composition. Fat mass, lean mass and total body weight were measured by MRI (*n*=8 per group). (D) Representative pictures of brown adipose tissue (BAT), inguinal white adipose tissue (iWAT) and liver. (E) The average tissue weights are presented as % of body weight (*n*=6 per group). Statistical comparison between groups was performed using Student’s *t* -test (* p < 0.05 and ** P < 0.01). All quantitative data for mice are presented as means ± SEM.

**Figure S2.**
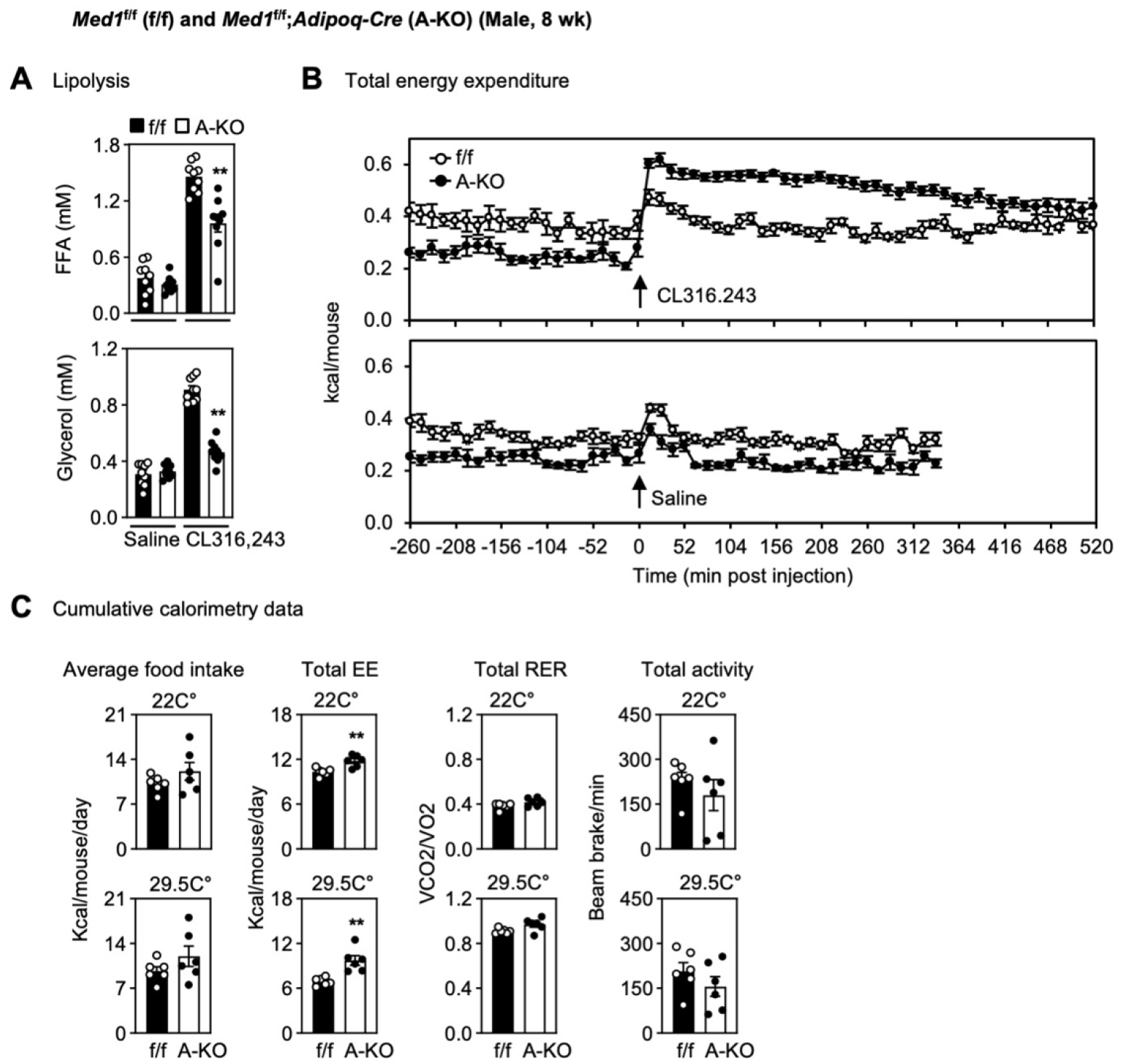
Related to Figure 1. Impaired energy expenditure in *Med1*^f/f^;*Adipoq-Cre* mice. 8wk male f/f and A-KO mice (*n*=8 per group) were fed with a regular diet. (A) Lipolysis analysis. Serum levels of FFA or glycerol were measured after saline or CL316,243 administration. (B-C) Total energy expenditure (B) after saline or CL316,243 administration and total cumulative calorimetry data (C) are shown. EE, energy expenditure; RER, respiratory exchange ratio. Statistical comparison between groups was performed using Student’s *t* -test (** P < 0.01).

**Figure S3.**
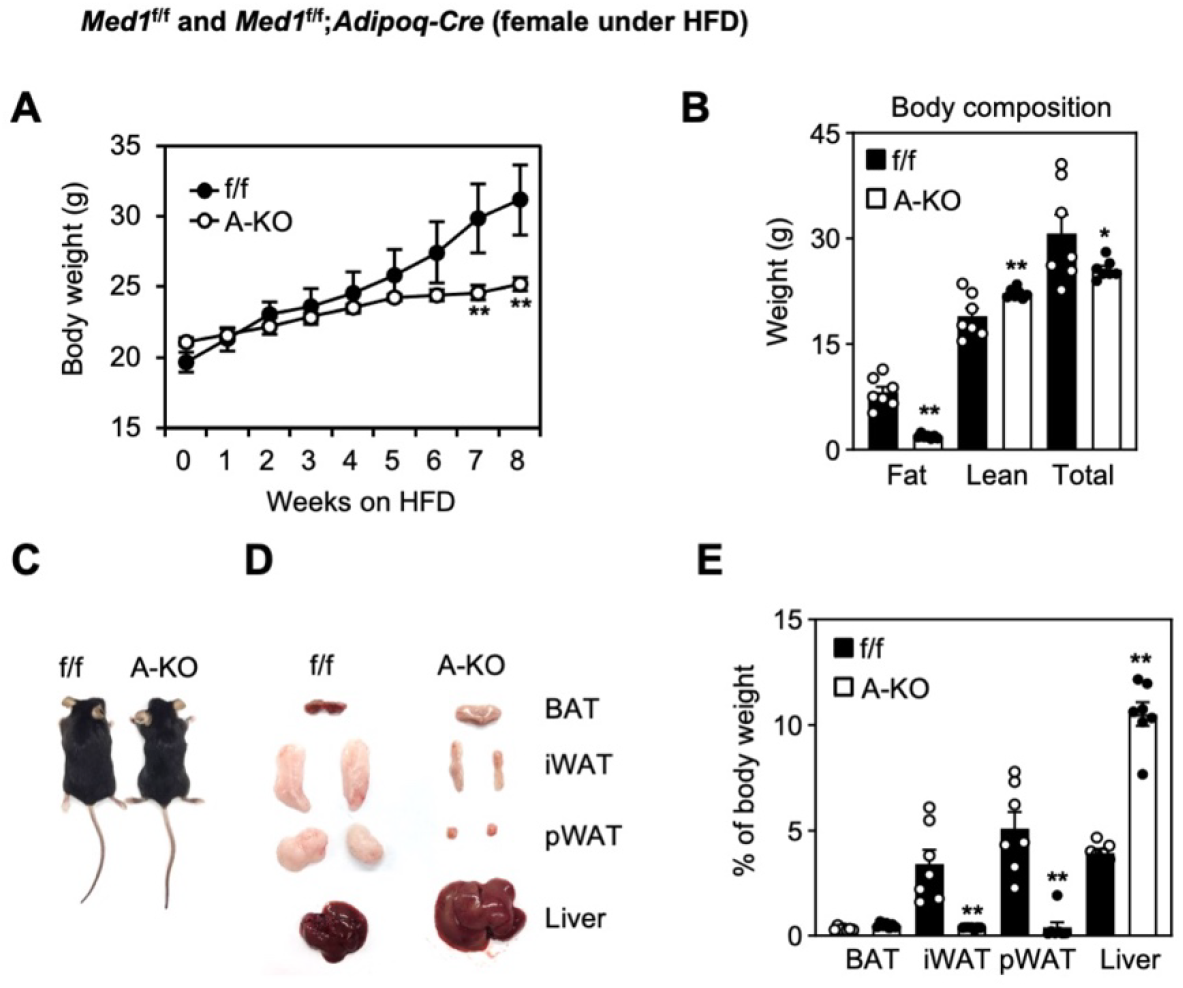
Related to Figure 2. Female mice with adipocyte-selective deletion of *Med1* are resistant to high fat diet-induced obesity. Female f/f and A-KO mice (*n*=7 per group) were fed with a high fat diet (HFD) from the 8th week of age. (A) Total body weight. (B) Body composition. Fat mass, lean mass and total body weight were measured by MRI. (C) Representative morphology of HFD-fed mice. (D) Representative pictures of BAT, iWAT, perigonadal WAT (pWAT) and liver. (E) The average tissue weights are presented as % of body weight. Statistical comparison between groups was performed using Student’s *t* -test (** P < 0.01 and * P < 0.05).

**Figure S4.**
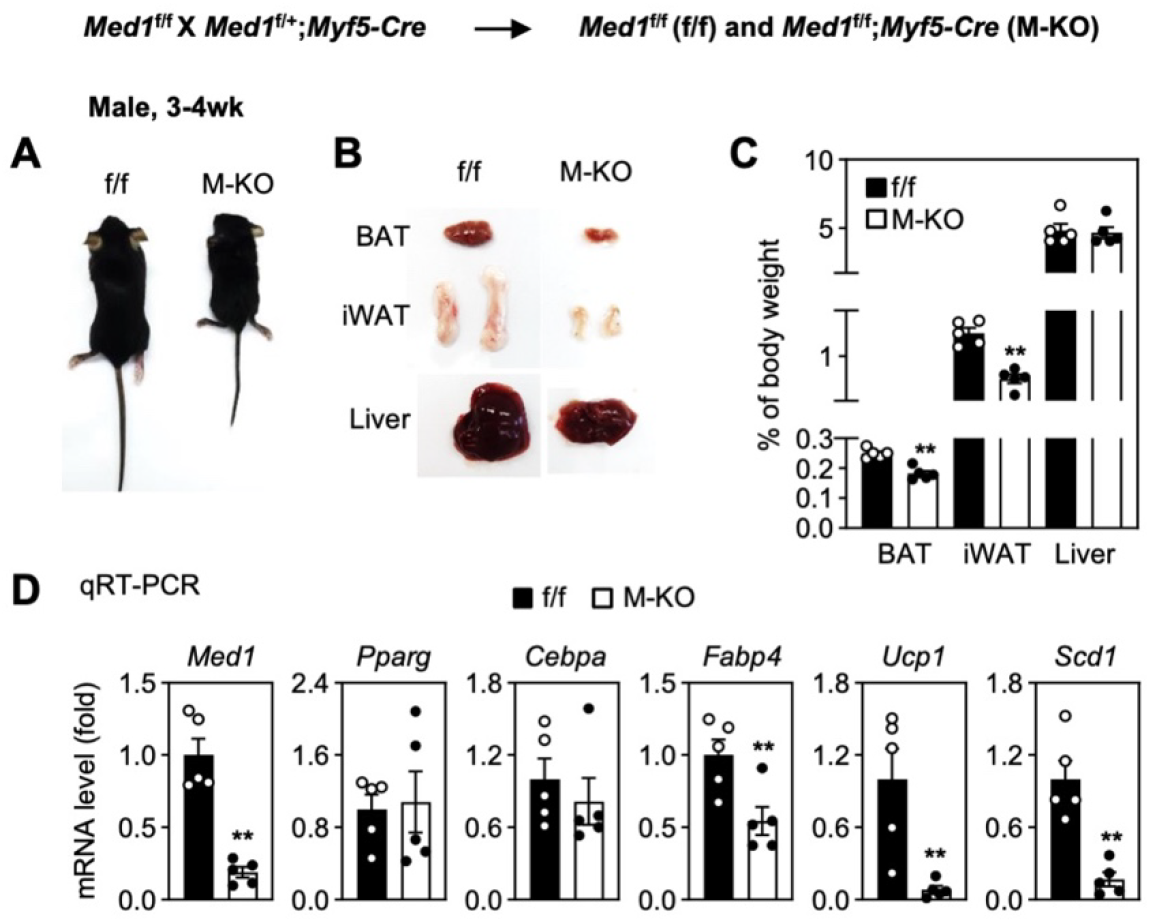
Related to Figure 3. *Med1*^f/f^;*Myf5-Cre* mice show growth retardation. *Med1*^f/f^ (f/f) mice were crossed with *Myf5-Cre* to generate *Med1^f/f^;Myf5-Cre* (M-KO) mice. (A) Representative morphology of 3-4wk males. (B) Representative pictures of BAT, iWAT and liver. (C) The average tissue weights are presented as % of body weight (*n*=5 per group). (D) qRT-PCR of *Med1, Pparg, Cebpa, Fabp4, Ucp1* and *Scd1* expression in BAT of mice at 3-4wk (*n*=5 per group). Statistical comparison between groups was performed using Student’s *t* -test (** P < 0.01).

**Figure S5.**
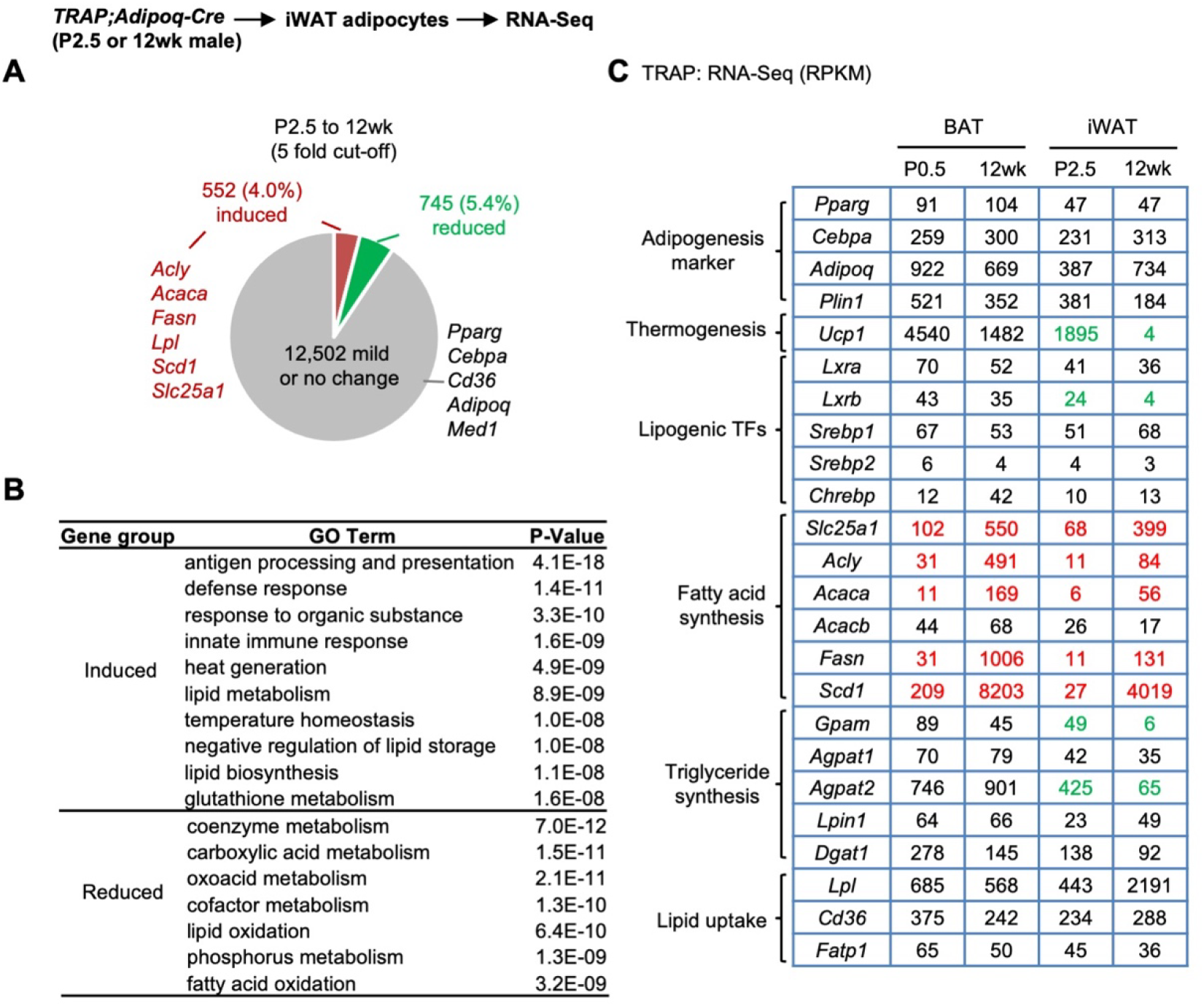
Related to Figure 4. Marked induction of lipogenesis genes in white adipocytes from newborn to adult mice. (A) RNA-Seq analysis of Adipoq^+^ white adipocytes isolated from iWAT of *TRAP;Adipoq-Cre* mice at P2.5 and 12wk. The cut-off for induced or reduced genes from P2.5 to 12wk is 5-fold. (B) GO analysis of gene groups defined in (A). (C) Expression levels of representative genes are shown in RPKM values of RNA-seq data from BAT and iWAT of *TRAR;Adipoq-Cre* mice. *Gpam* and *Fatp1* are also known as *Gpat1* and *Slc27a1*, respectively.

**Figure S6.**
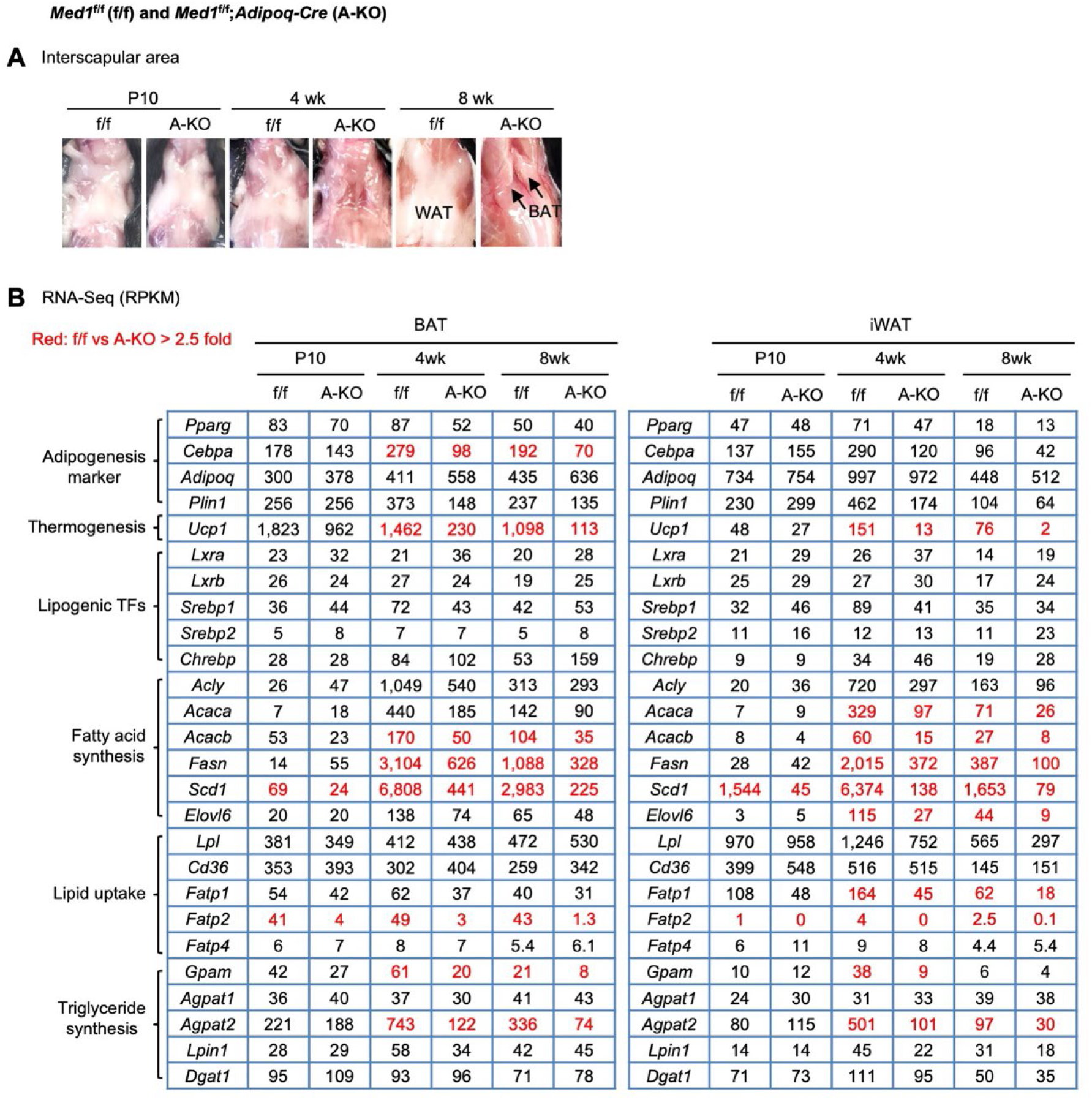
Related to Figure 4. MED1 is required for postnatal adipose expansion and lipogenesis gene expression. BAT and iWAT were collected from f/f and A-KO mice at P10, 4wk and 8wk. Total RNA isolated from 3-5 mice per genotype was combined in equal amounts for RNA-Seq. (A) Representative morphology of interscapular area at P10, 4wk and 8wk. (B) Expression levels of representative genes are shown in RPKM values.

**Figure S7.**
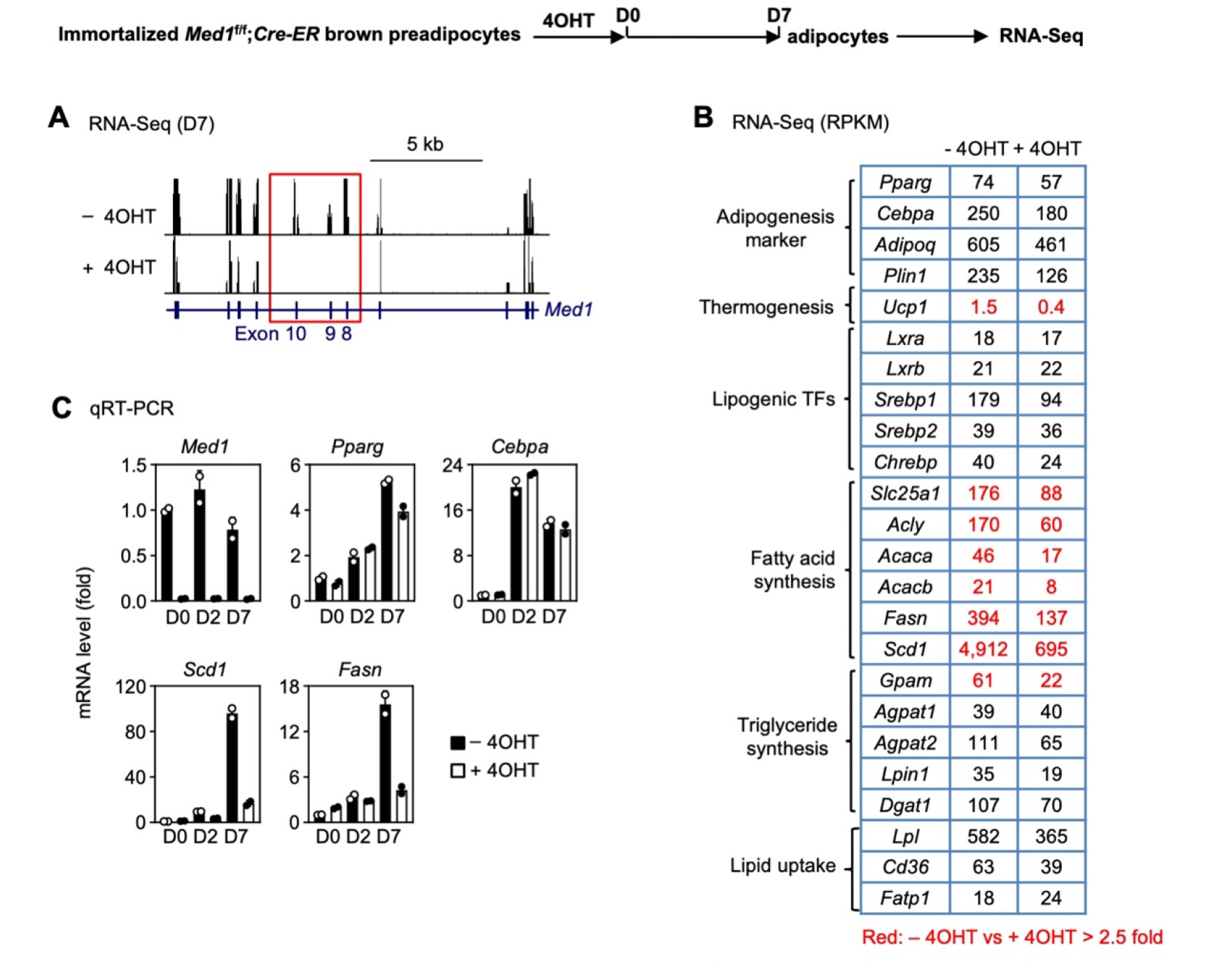
Related to Figure 5. MED1 is required for lipogenesis but not early adipogenesis in brown adipocytes. Adipogenesis was done as in Figure 5, followed by RNA-Seq and qRT-PCR. (A) Profiles of RNA-Seq data on *Med1* gene locus. The targeted exons 8-10 are highlighted in red box. (B) Expression levels of representative genes are shown in RPKM values. (C) qRT-PCR of *Med1, Pparg, Cebpa, Scd1,* and *Fasn* expression at D0, D2 and D7 of adipogenesis. All qPCR data in cells presented as means ±SD. Two technical replicates from a single experiment were used.

**Figure S8.**
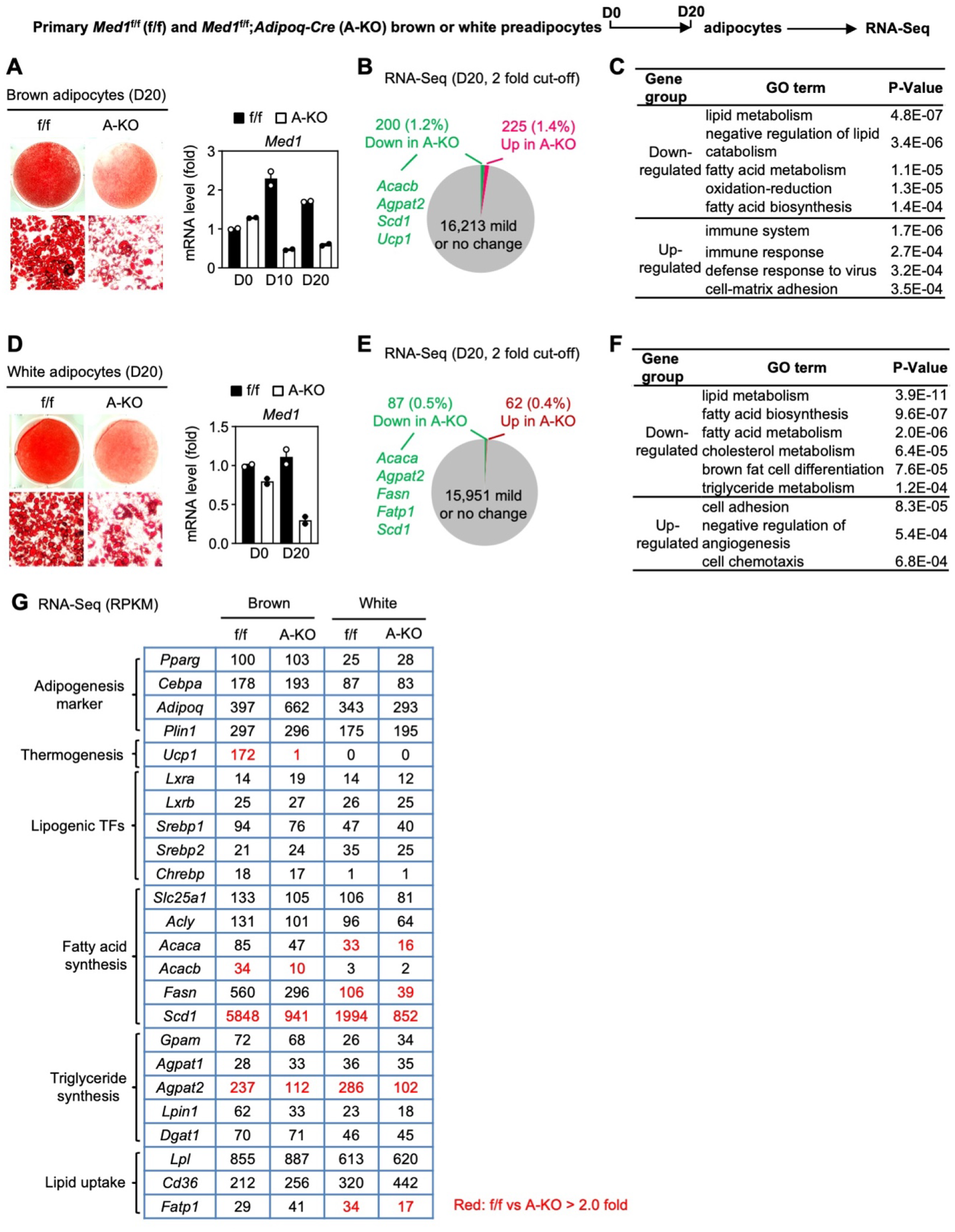
Related to Figure 5. Deletion of *Med1* reduces lipogenesis gene expression in primary brown and white adipocytes. Primary brown and white preadipocytes were isolated and cultured from f/f or A-KO mice, followed by adipogenesis assay and RNA-Seq. White preadipocytes were treated with Rosiglitazone during adipogenesis for more efficient adipogenesis. (A, D) Oil Red O staining at D20 of adipogenesis (left panel) and qRT-PCR of Med1 in mature adipocytes (right panel) are shown for brown (A) and inguinal white adipocytes (D). (B, E) Pie charts show the identification of down- or up-regulated genes in A-KO brown (B) and white adipocytes (E) at D20. The cut-off for differential expression is 2-fold. (C, F) GO analyses of gene groups defined in (B, E) are shown. (G) Expression levels of representative genes are shown in RPKM values

**Figure S9.**
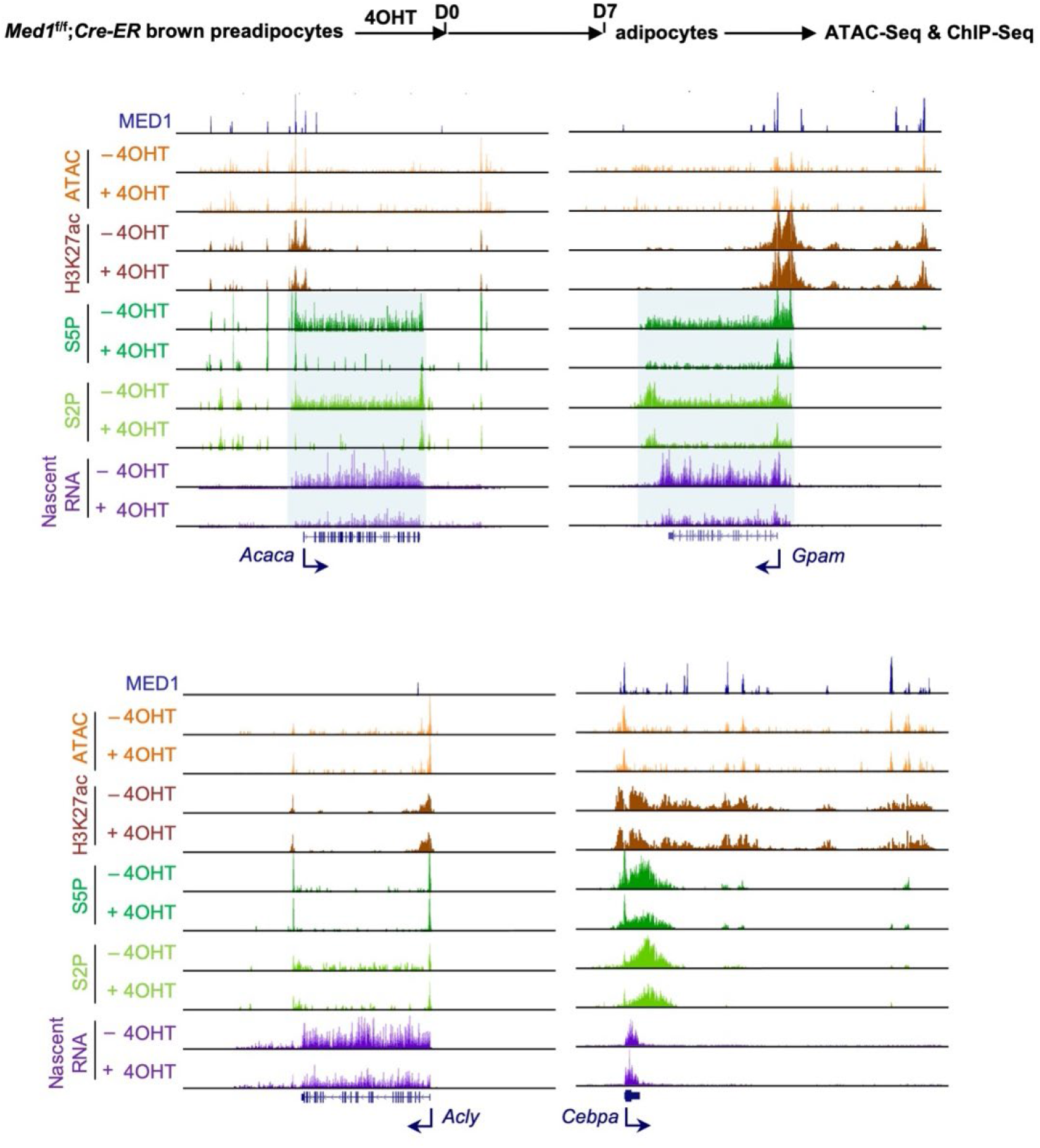
Related to Figure 6. MED1 is required for Pol II binding on lipogenesis genes in adipocytes. *Med1^f/f^;Cre-ER* brown preadipocytes were described in Figure 6. Profiles of ATAC-Seq, H3K27ac enrichment, S5P-Pol II, S2P-Pol II binding, and nascent RNA-Seq data on *Acaca*, *Gpam, Acly* and *Cebpa* gene loci are shown.

**Figure S10.**
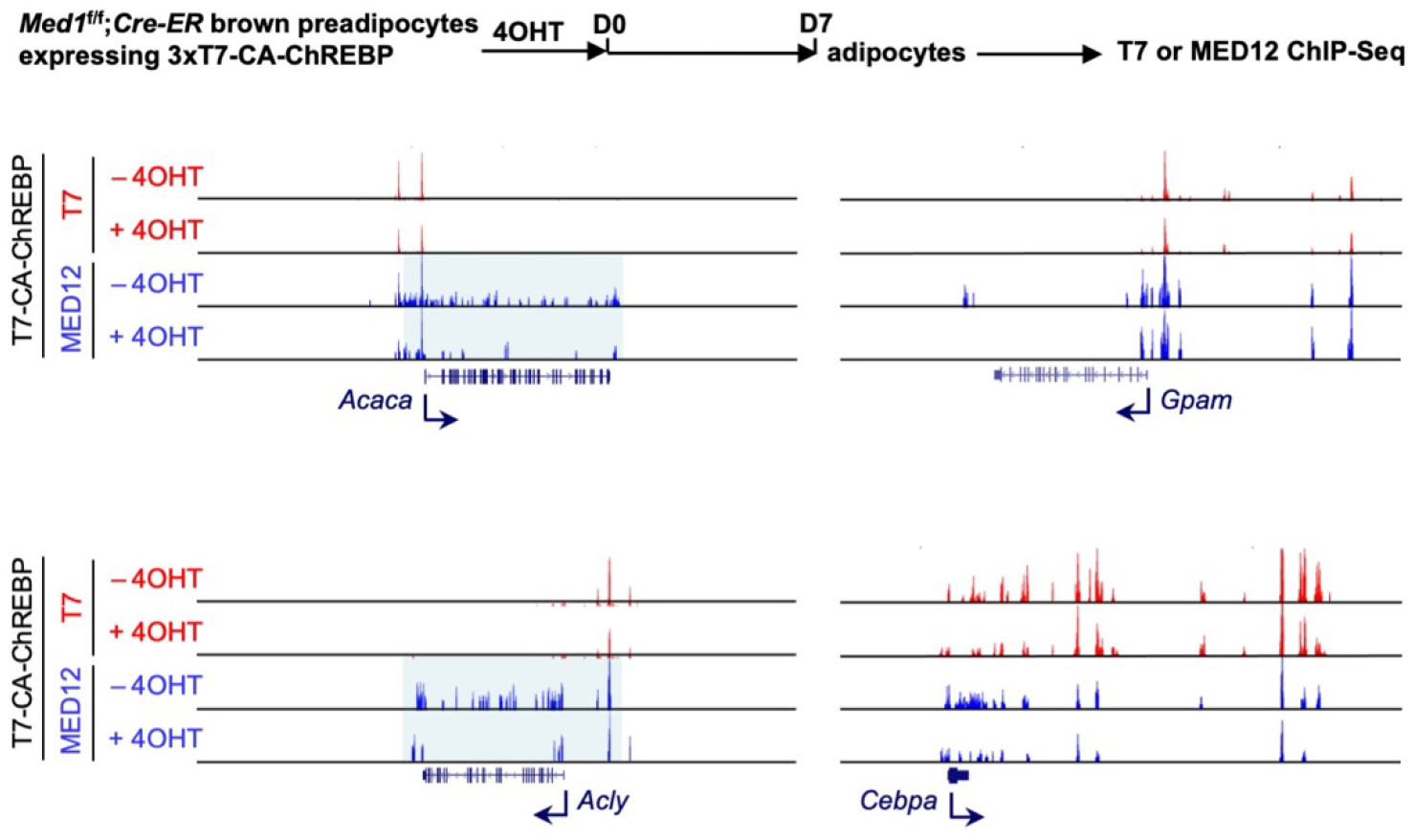
Related to Figure 7. MED1 is required for Mediator binding on a subset of lipogenic enhancers. *Med1*^f/f^;*Cre-ER* brown preadipocytes expressing 3xT7-CA-ChREBP were described in Figure 7. Profiles of T7-CA-ChREBP and MED12 binding on *Acaca, Gpam, Acly* and *Cebpa* gene loci are shown.

## Notes

### Competing Interest Statement

The authors have declared no competing interest.

